# Continuous estimation of reaching space in superficial layers of the motor cortex

**DOI:** 10.1101/2023.12.01.569527

**Authors:** Gregorio Luis Galiñanes, Daniel Huber

## Abstract

Motor cortex plays a key role in controlling voluntary arm movements towards spatial targets. The cortical representation of spatial information has been extensively studied and was found to range from combinations of muscle synergies to cognitive maps of locations in space. How such abstract representations of target space evolve during a behavior, how they integrate with other behavioral features and what role they play in movement control is less clear. Here we addressed these questions by recording the activity of layer 2/3 (L2/3) neurons in the motor cortex using two-photon calcium imaging in head-restrained mice, while they reached for water droplets presented at different spatial locations around their snout. Our results reveal that a majority (>80%) of L2/3 neurons with task-related activity are target-space selective and their activity is contingent on a single target position in an ego-centric reference frame. This spatial framework is preferentially organized along three cardinal directions (Center, Left and Right). Surprisingly, the coding of target space is not limited to the activity during movement planning or execution, but is also predominant during preceding and subsequent phases of the task, and even persists beyond water consumption. More importantly, target specificity is independent of the movement kinematics and is immediately updated when the target is moved to a new position. Our findings suggest that, rather than descending motor commands, the ensemble of L2/3 neurons in the motor cortex conjointly encode internal (behavioral) and external (spatial) aspects of the task, playing a role in higher-order representations related to estimation processes of the ongoing actions.

## INTRODUCTION

Manipulating objects in our surroundings is an integral part of our everyday life and is key to survival. Rodents, much like primates, rely on their hands to manipulate and eat food and deploy motor programs specially adapted to each type of food ^1,2,3^. Coordination and execution of such complex behaviors occur at multiple levels of the neuraxis, with the motor cortex playing a central role ^4,5^. For instance, microstimulation, inactivation, anatomical and physiological experiments suggest that the medial part of the anterior motor cortex (Medial Anterior Cortex, MAC ^6^) is involved in controlling reach-to-grasp movements in the mouse ^7,3,8^. Moreover, there is convergent evidence across different species that the cerebral cortex is functionally organized into distributed action maps that orchestrate ethologically relevant behaviors ^9–12^. Although the exact mechanisms by which the motor cortex controls movement are still debated^13^, it is generally accepted that cortical output is generated in layer 5b as the result of local dynamics and inputs from other cortical areas and the thalamus ^14,15^. While recent studies have started to elucidate the role of thalamocortical inputs in motor preparation, movement timing and activity pattern generation ^16,17,14^, the origin and nature of additional external inputs is less clear ^16^.

Most motor control theories emphasize that the optimal generation of motor commands is achieved through the continuous integration of feedback signals to guide movement ^18–21^. The brain uses sensory information, such as proprioception and vision, to estimate the current state of the body and environment, which, combined with the current motor goal, helps in computing the best motor commands to achieve a successful movement^22^. Recent evidence suggests that such state estimation mechanisms might take place in the parietal cortex and the cerebellar circuits ^23,24^. However, whether and how these mechanisms influence motor cortex activity is not clear. From a cytoarchitectonic point of view, layer 2/3 neurons are optimally connected to fulfill this role, since they receive multimodal inputs from other cortical areas and they are hierarchically connected to layer 5b neurons ^15,25^.

To investigate a possible role of layer 2/3 motor cortex neurons in state estimation mechanisms, we decided to utilize a behavioral paradigm for head-fixed mice, inspired in the center-out reaching task of primates^8^. In this task, water-deprived mice perform directional reaching movements to retrieve a water drop from a waterspout that entered and exited the peripersonal space of the mouse at different locations in front of their snout. The dynamics of this paradigm created opportunities for specific motor actions depending on the current state of the animal and its spatial relation to the object, thereby providing complex and naturalistic real-world-like interactions. By performing simultaneous two-photon imaging recordings, we found that layer 2/3 neurons of the MAC are simultaneously dictated by the location of the waterspout and by the ongoing action of the animal. This joint representation of space and behavior is expressed across all phases of the task and reflects a general role of continuous integration of internal and external feedback, compatible with state estimation mechanisms.

## RESULTS

### Reaching maps in layer 2/3 of the MAC

Neurons in the primate motor cortex are widely recognized for the encoding of kinematic parameters of movement. This is particularly evident in the center-out reaching task, where population coding of arm directionality has been thought to be one of its defining characteristics ^26^. It is only recently, however, that studies have begun to investigate how directional arm movements are encoded in the rodent cortex ^8,27,28^. Through anatomical, optogenetic and microstimulation experiments, the MAC has been pinpointed as the principal cortical region related to the control of reaching in mice ^6,3,8^. Therefore, we aimed to determine the tuning properties of reach-related neurons within this region by performing two-photon microscopy imaging of layer 2/3 neurons while head-fixed mice reached for drops of water presented at 3 different locations around their snout (Left, Center and Right, Figure 1ab, supplementary figure 1).

We first focused our analysis on reach-related neurons, as defined by responding within a 500 ms time window before reach onset and up until the first touch of the waterspout. Reach-related neurons were readily observed across layer 2/3 of the MAC with an average response time of −53±18.9s (mean ± S.D.) from reach onset. Over half of the reach-related neurons (57.4%) responded before movement initiation and the rest responded during the reaching movement.

As previously reported ^8^, a striking feature of neurons in MAC is their pronounced trial type modulation, with a discernible lateralization bias towards the right and individual neurons distinctly active depending on whether the mouse reached to the left, center, or right. (Figure 1C, Supplementary figure 2). This type of activity is reminiscent of directional neurons in the primate motor cortex. However, reach-related neurons in the mouse appeared to display an ‘all or nothing’ response pattern, rather than the cosine tuning pattern observed in primates ^29,30^. Indeed, the majority of the neurons (81.3±2.0%) were selectively active when the animal reached towards only one of the three target locations (Figure 1c,d).

**Figure 1:**
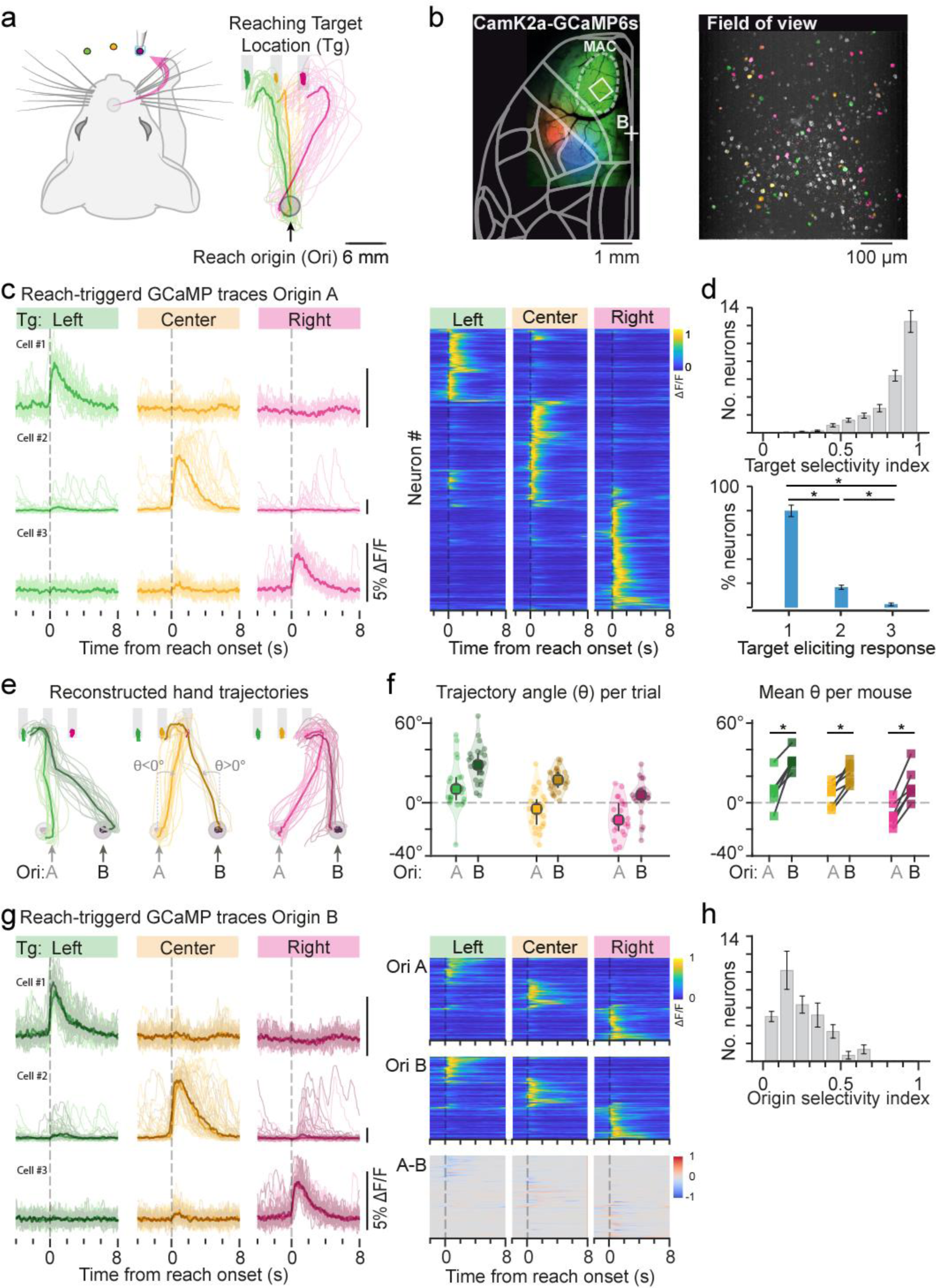
Layer 2/3 MAC neurons encode reaching target location **a** Left: schematic representation of the three-target water reaching task where head-fixed mice reached for, retrieved and drank water droplets with their right hand from multiple locations around their snout. Reaching movements initiated at the position determined by the resting bar (Reach origin) and were performed towards a waterspout the position of which was determined on a trial by trial basis using a motorized system. Right: frontal view (mirrored) hand reconstructed trajectories of one example session. **b** Cranial window on the left hemisphere of a mouse expressing GCaMP6s in excitatory neurons (green) providing optical access to the primary and secondary motor cortices as well as the upper limb (red) and lower limb (blue) somatosensory cortical regions. Somatosensory areas are shown in pseudocolor after wide field somatosensory mapping. MAC: medial anterior cortex. Square: field of view (FOV) during two-photon excitation microscopy (right). **c** Left: Perievent calcium traces (PECT) of three reach-related neurons during Left, Center, and Right trials. Thin lines individual trials, thick line median average. Neurons display a strong selectivity for the target location. Right: Normalized average activity of all reach-related neurons (1226 neurons, 13 mice, 44 FOVs). **d** Top: Distribution of the Target Selectivity Index for all reach-related neurons. Zero indicates equal response amplitude for all target locations; 1 indicates null response amplitude for two of the targets. Bottom: 81.3±2.0% of the neurons significantly responded to only one reaching location, 16.1±2.6% to two, and 2.66±0.9% to all three. **e** Reconstructed hand trajectories of an example session to the Left, Center and Right target location when reaching initiated from two different origins (Ori A, Ori B). **f** Left: Trajectory angles (θ) defined as the angle between the hand trajectory and a line perpendicular to the ground. Small dots individual trajectories shown in (e). Right: Session average trajectory angles per mouse (6 mice). **g** Left: PECTs (ΔF/F) of the same cells in (c) during reaching trials from Ori A (lighter colors) and Ori B (darker colors). Right: normalized average responses of all recorded neurons during reaching trials from Ori A (top), Ori B (middle) and subtracted responses A-B (bottom, 192 neurons, 5 mice, 6 FOVs). **h** Origin Selectivity Index for all neurons shown in (g). Zero indicates equal response amplitude to both reaching origins; 1 indicates null response amplitude for one of the reaching origins. *: p<0.01. Bars: mean; error bars: S.E.M

This response pattern raised the possibility that reach-related activity could be associated with reaching target locations, rather than the movement direction of the arm ^31^. To address this question, we analyzed the activity of the same neurons while the animal initiated reaching from two different origins (Figure 1e,f). This manipulation forced the animals to execute different arm trajectories to reach the same endpoint, thereby decoupling both factors (Figure 1e). Contrary to our expectation, the neuronal activity remained unaffected by this manipulation (Figure 1g-h) suggesting that these neurons are dictated by the endpoint but not by the starting point or the trajectory of reaching. Importantly, the activity of these neurons was specific to which arm performed the task (Supplementary figure 3), underscoring that reach-related neuronal activity in the MAC is actually linked to the execution of hand movements. Furthermore, these results highlight that a prominent feature encoded by layer 2/3 neurons is the location of the reaching target and suggest that the MAC encodes an egocentric reaching endpoint map instead of a movement kinematics map.

### Properties of cortical reaching maps

To more precisely quantify the spatial representation of the reaching space, we increased the number of target locations from three to seven (Figure 2a). Consistent with our initial findings, reach-related neurons exhibited a strong preference for specific target locations, as evidenced by a high target selectivity index (Figure 2b-d). Interestingly, the finer mapping of the reaching space revealed that reach-related neurons possess small response fields of an average width of 4.42 ± 2.02 mm on the mediolateral axis (supplementary figure 4). As a consequence of a smaller spacing between target locations, in the seven-target variant of the task, the majority of the neurons responded to up to 3 contiguous reaching locations (Figure 2e).

**Figure 2:**
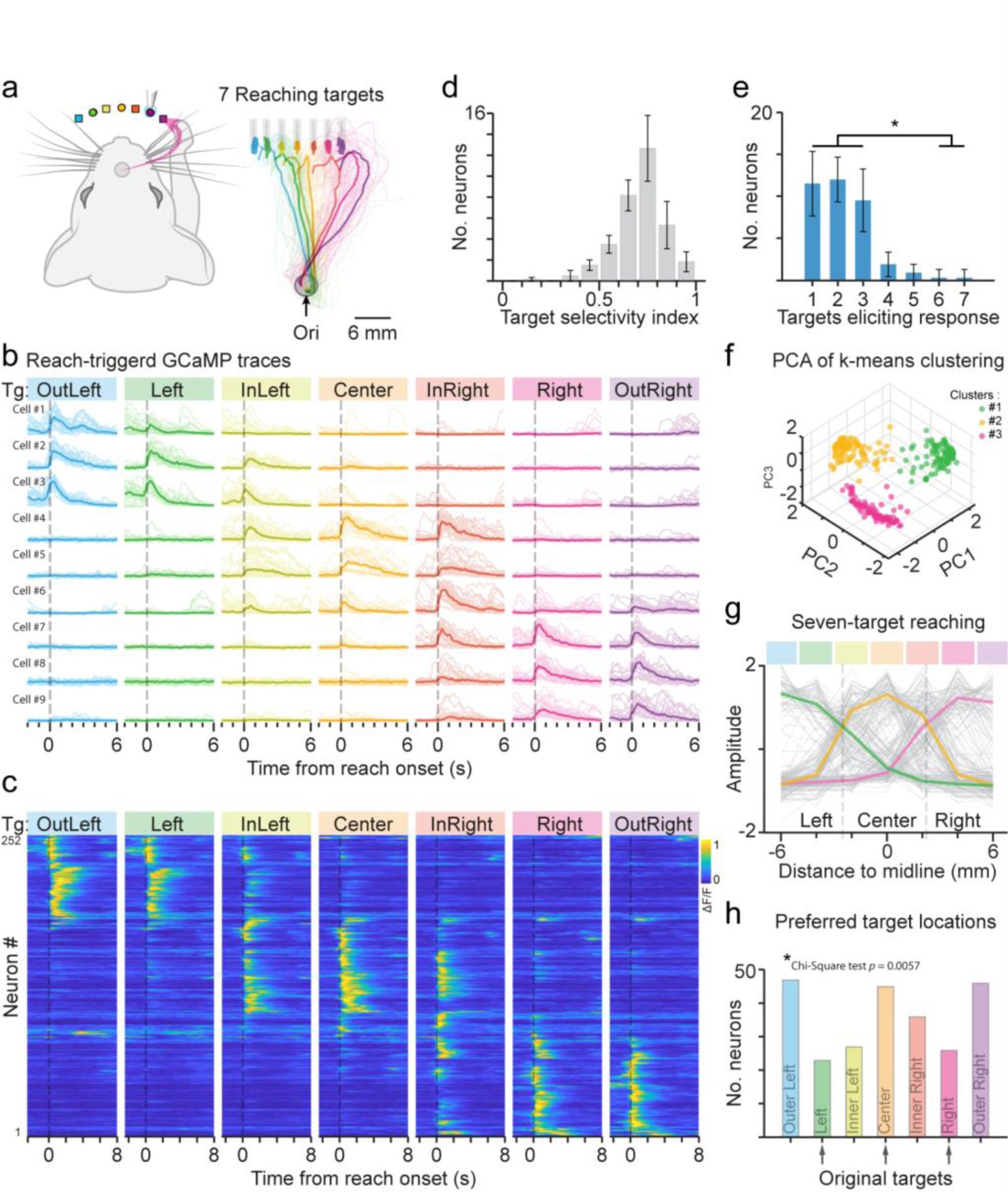
Reaching target space is divided into three main areas. **a** Left: schematic representation of the seven-target reaching task. Original targets (Left, Center, Right) are represented with a circle. Novel targets (Outer-Left, Inner-Left, Inner-Right, Outer-Right) with a square. Right: frontal view (mirrored) hand reconstructed trajectories. **b** PECTs (ΔF/F) of nine representative reach-related neurons aligned to reach onset during the seven-target reaching task. **c** Normalized average activity of all reach-related neurons (298 neurons, 6 mice). **d** Distribution of the Target Selectivity Index for all reach-related neurons in the seven-target reaching task. Zero indicates equal response amplitude for all target locations; 1 indicates null response amplitude for six of the targets. **e** Most of the neurons (91.6±2.6%) responded to less than 3 reaching targets. **f** A 3D representation of k-means clusters of the seven-target response profiles. Each data point represents a neuron with a color code according to its response bias. **g** Z-scored response amplitude profile of individual neurons (n=250, gray lines) across the mediolateral axis reveals the boundaries (dashed lines) between the three main response fields of the reaching target space. Colored lines: Average response profile for each of the k-means clusters color coded according to their spatial preference. **h** Histogram of neurons’ maximal responses to each of the seven target locations. Chi-square test showed significant deviation from the uniform distribution, with a noticeable bias towards Center, Outer-Left, and Outer-Right locations. Bars: mean; error bars: S.E.M. * p<0.015

At the population level, the distribution of response fields did not form a continuous map of the reaching target space (Figure 2b,c). Instead, the spatial selectivity of cortical neurons gave rise to three distinct clusters, each of which showed a preference for target locations around the left, center, or right of the animal (Figure 2f,g, supplementary figure 5). This indicates a compartmentalized spatial representation, demarcated by relatively sharp boundaries (Figure 2c,g). Notably, these boundaries were observed at the level of some individual neurons (Supplementary Figure 4), which responded robustly within their preferred area of the reaching space but exhibited no response for the immediate neighboring locations. Interestingly, the division of the reaching space into three cardinal directions is biased towards the Center, Outer-Left and Outer-Right positions (Figure 2h), and does not appear to be the result of training. Recordings from naïve animals performing the three-target reaching task for the first time revealed the same spatial compartmentalization (Supplementary figure 5). Taken together, the data suggest a potential innate role for MAC neurons as spatial location classifiers.

### Encoding of task events beyond reaching

The majority of studies of motor control typically focus on a range of behavioral phases strictly related to movement, such as instruction, preparation and execution. However, neurons in motor and premotor cortices have been implicated in higher order phenomena that extend beyond movement, such as reward processing, behavioral outcome or memory traces ^32,33,34^. Whether neurons in the MAC encode these kinds of signals and how spatial information is integrated with them, is less clear. To answer these questions, we took advantage of the multiple phases of the water reaching task (Supplementary figure 1) and investigated the neuronal activity of the MAC beyond reaching.

We identified a substantial number of neurons (57.2% of all task-related neurons) that were active during different phases of the task, from the start to the end of the trial and during the inter-trial interval. Besides reach-related neurons, the second largest proportion of neurons responded at trial start and during the delay period, followed by neurons active during water consumption, and finally, by neurons active during the inter-trial period. Purely touch-related neurons were also observed, but in many occasions their activity extended to neighboring phases. Indeed, many neurons, including the reach-related ones, displayed persistent activity and were active during more than one phase of the task, with a few extreme cases where the activity spanned from waterspout presentation to waterspout retraction (Supplementary figure 6).

When we analyzed the spatial tuning properties of all task-related neurons, we found that they, too, were modulated by the location of the reaching target. Surprisingly, this modulation was as pronounced as that of reach-related neurons (supplementary figure 7), suggesting that task-related neurons may collectively form functional sets determined by the location of the reaching target. Indeed, this tuning was so prominent that it was already visible at the level of spontaneous activity of task-related neurons and in the pairwise cross-correlation matrix (figure 3a,b).

**Figure 3:**
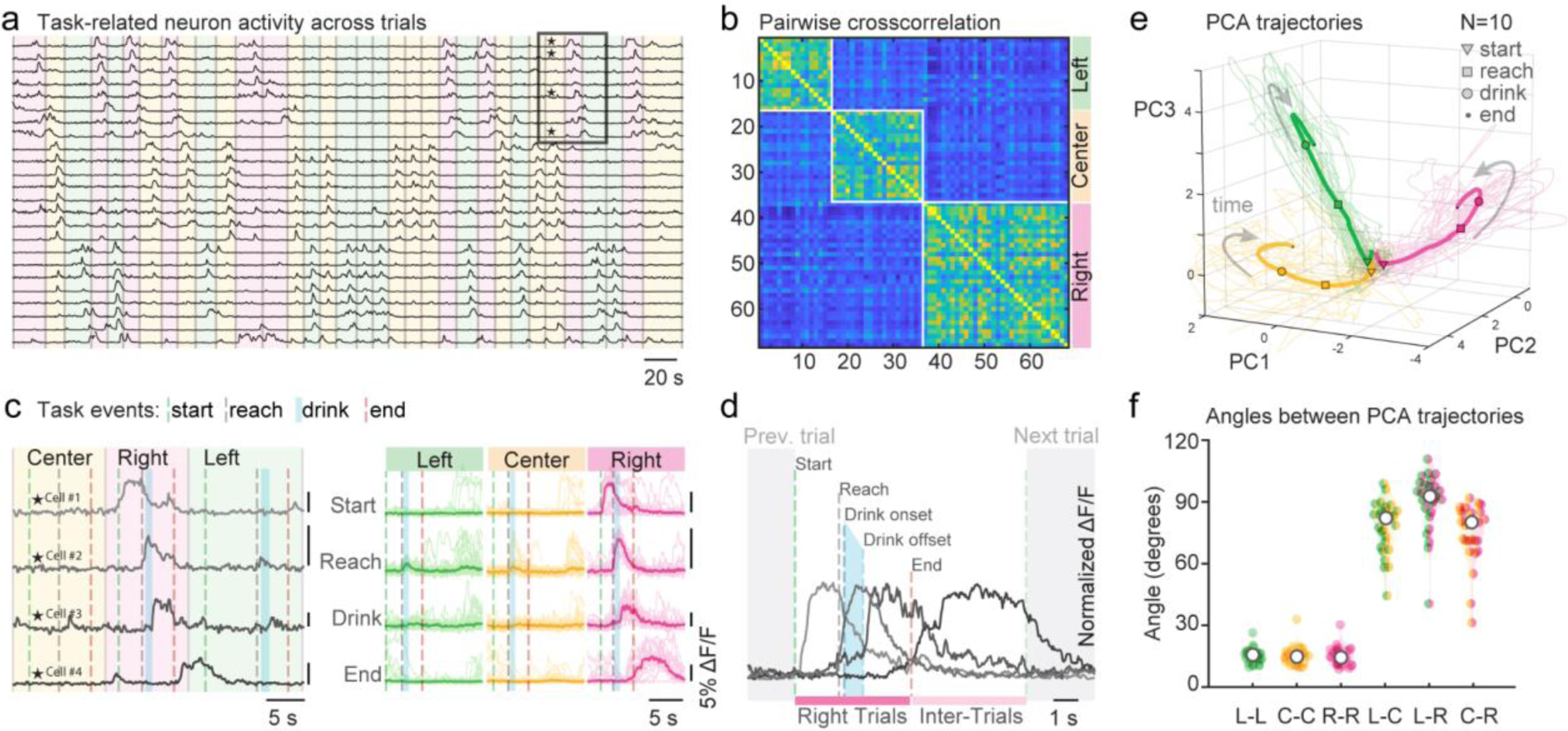
Task events are conjointly represented with the reaching target location. **a** Normalized ΔF/F of task related neurons (subset) during a three-target reaching session. Green, yellow, and pink shades indicate the reaching target location for each trial (Left, Center and Right). Neurons were sorted according to their trial type preference. **b** Pairwise cross-correlation matrix of all task-related neurons (n=68) recorded during the session in panel b reveals the presence of three clusters, each corresponding to a reaching target location. **c** Left: Zoom in of three consecutive trials (gray square in (a)) showing four task-related neurons sequentially active across different phases of the task. Task events s:start (target presentation), r:reach, d:drink and e:end (target retraction). Right: PECTs (ΔF/F) of the same neurons pseudo-aligned to task-event triggers according to the reaching target location. Thin traces, individual trials; thick trace, median average. Vertical lines indicate the average reference time of the triggering task events. **d** Average activity of the four neurons shown in (c) pseudo-aligned to the average task event timings (dashed lines). **e** Neural trajectories of dimensionally reduced data (PCA) from trial start to trial end based on the reaching target location. Thin lines represent the average activity from individual sessions (10 mice, 27 FOVs). Thick lines indicate the mean across all sessions. **f** Pairwise angles between session-averaged PCA trajectories (shown in e) for trial types (L: Left, C: Center, R: Right; 10 mice, 27 FOVs)

The onset and offset boundaries of single neurons’ activity were generally sharp and determined by specific external events (e.g. target presentation and retraction), or mouse’s actions (e.g. drink beginning and ending) in conjunction with the reaching target location. Therefore, neurons with the same spatial tuning revealed a sequential and continuous representation of all the phases of the task as a chain of events (figure 3c,d). We confirmed this finding by applying a dimensionality reduction technique of the full dataset (principal component analysis, PCA, supplementary figure 8), where we observed that the neural state space trajectories diverged upon waterspout presentation and remained separated across all the phases of the task beyond target retraction. These trajectories followed near-orthogonal paths, each determined by the location of the reaching target (figure 3e,f). Importantly, PCA of neurons that were classified as non-task-related, revealed that they did not carry significant information related to the task (supplementary figure 8), suggesting that the organizing principle of cortical activity during the execution of this task is the location of the reaching target.

### Dynamic representation of task events

The data so far suggest that task progression in the MAC is represented by a sequence of task-related neurons whose activity is conjointly determined by the phase of the task and the location of the reaching target. If this was the case, sudden changes in the waterspout position should produce predictable changes in cortical activity. To test this, we designed a variant of the three-target reaching task where, immediately after reach onset, the waterspout was unexpectedly relocated (“jumped”) from one target location (“Source”) to another (“Destination”). Mice readily performed this variant of the task executing hand movement corrections towards the new position of the waterspout (Figure 4a, Supplementary Figure 9).

**Figure 4:**
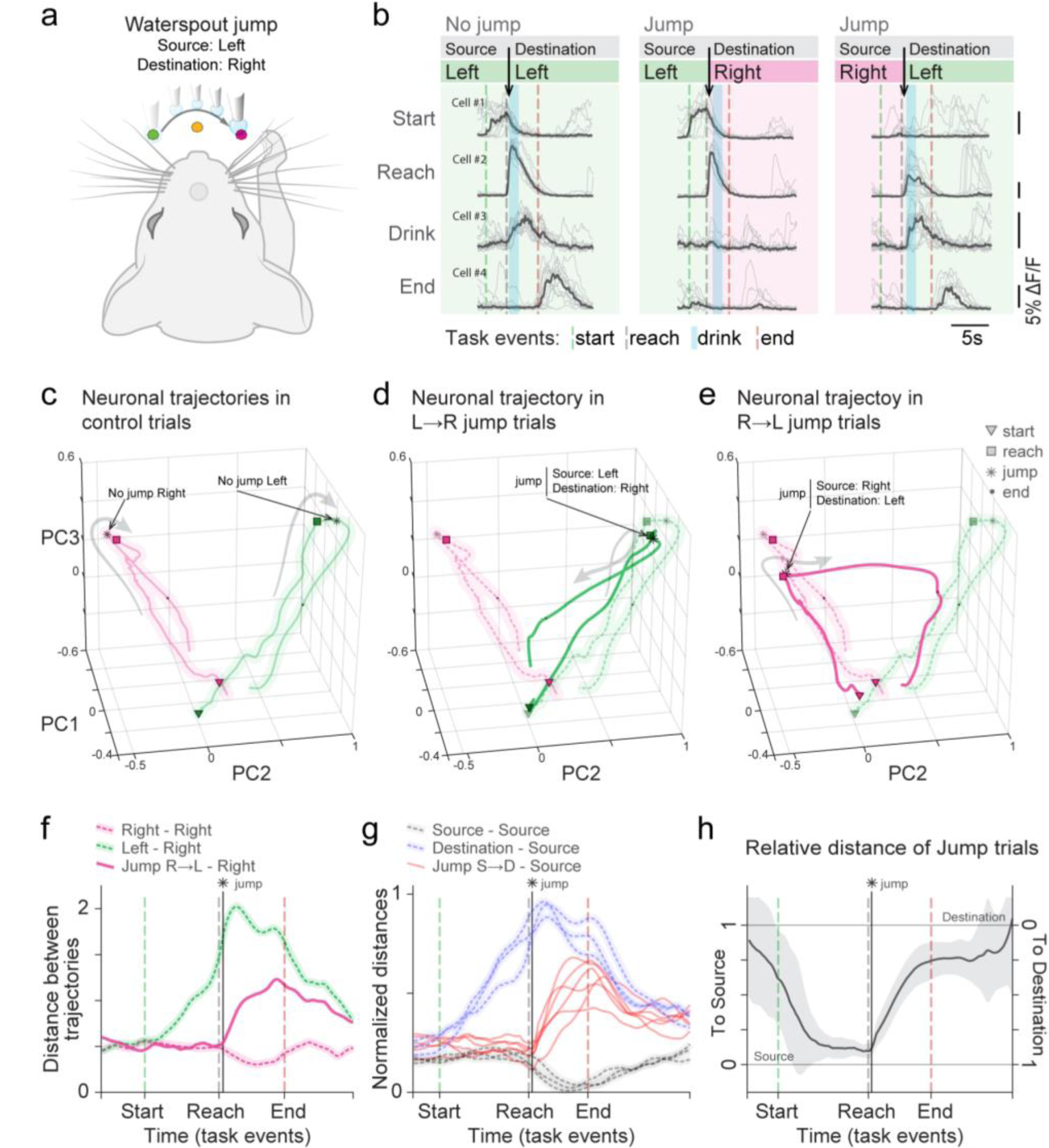
Cortical neurons dynamically represent ongoing task events and target locations. **a** Schematic representation of the waterspout jump task. In a subset of randomly selected trials the waterspout jumps to another target location (e.g. from Left to Right) after reaching initiation. **b** Pseudo-aligned PECTs of four sequentially active neurons during different phases of the task in control (“no-jump”) and jump trials (Left jumps to Right and Right jumps to Left). Black arrow indicates the moment of waterspout jump and color shades the target location (green: Left; pink: Right). Task events are indicated by dashed lines: start (target presentation), reach onset, drinking period and end (target retraction). **c** Average trajectories in the neural state space from neurons recorded simultaneously with those in (b), during control (no-jump) trials for Left and Right waterspout locations. Gray arrows denote time progression. **d-e** Extensions of (c) but including jump trials: **d** Trajectory with waterspout source at Left and jump destination at Right. **e** Trajectory with source at Right and destination at Left. **f** Average bootstrapped distances between neural state trajectories shown in (e) across task phases. Distances Left, Jump R→L and Right trials were computed against Right control trials. **g** Extension of (f) showing normalized distances for all source and destination combinations averaged across mice. **h** Relative distance of jump trajectories to control source and destination trajectories show that upon waterspout jump, the neural state transitions away from the source approaching the destination condition. Bootstrapped of average trajectories from all source and destination combinations (3 mice, 4 FOVs).

The leftmost panel of figure 4b shows the activity of four example neurons that were sequentially active at different phases of control trials with the waterspout on the Left. The first two neurons were active during the “start” and “reach” phases, while the remaining two, were active during the “drink” and “end” phases of the task (Figure 4b “no jump” trials). The middle panel in figure 4b shows the activity of the same neurons during trials in which the waterspout jumped from the Left (source) towards the Right (destination) locations. As expected, the activity of the “start” and “reach” neurons (Cell #1 and #2) was not affected during these trials, because the waterspout was in their preferred location. Just after reach onset, the waterspout jumped to the Right location leading to the silencing of the “drink” and “end” neurons (Cell # 3 and #4). Finally, in the complementary jump trials (source on the Right and destination on the Left) the effect was reversed, meaning that the “start” neuron was not reactivated, but the “reach”, “drink” and “end” neurons were recruited once the waterspout was in their preferred location (Figure 4b, rightmost panel).

To determine the effects of this manipulation at the population level, we transformed the recorded activity to its PCs and compared the neural state trajectories of control and jump trials. As previously shown (Figure 3e), Left and Right control trials occupied near orthogonal regions of the neural state space (Figure 4c). On the other hand, jump trials initially followed trajectories similar to the control trajectories of the matching source target location, but upon waterspout jump, they diverted towards the neural state space corresponding to the destination target location (Figure 4d-h). Therefore, according to our prediction, the activity of MAC neurons was updated in real time to match the current location of the waterspout and the ongoing phase of the task. It is noteworthy that in addition to this, we observed the presence of neurons that were not active during control trials but were recruited exclusively during specific combinations of Source-Destination jump trials, pointing, as well, to a possible role in predictive processing mechanisms (Supplementary figure 9 ^35^)

Finally, we observed that the distance between neuronal trajectories of congruent trials (Source-Source distances, Figure 4g) consistently decreased after reach onset, throughout water drinking and during the intertrial period, suggesting a reduced neural state variability. This reduction in variability may be due to the accumulation of direct proprioceptive and tactile feedback information, leading to an increased level of certainty at the neuronal level. Taken together, our data indicate that the main role of L2/3 neurons in the MAC is to represent the ongoing behavioral state of the mouse in conjunction with the location of the reaching target suggestive of a higher-order function in behavioral control.

## DISCUSSION

Contrary to the traditional belief that the motor cortex encodes kinematic parameters, such as movement directionality ^36,26,29,37^, our data in mice provides a somewhat different picture. We demonstrate that during the water reaching task the principal determinant of neuronal activity within layer 2/3 of the MAC is, in fact, the spatial location of the target. This spatial influence unexpectedly extends well beyond the execution of target-oriented movements, encompassing all the phases of the task, even the intertrial period when the target has been retracted. Thus, more than movement features, the MAC exhibits an overarching encoding capacity, continuously representing the ongoing behavior of the mouse in relation to the most recent location of the reaching target.

Interestingly, the neuronal tuning patterns and spatial coding in the MAC bear stronger similarities to the activity in the PMd than that of the primary motor cortex of primates ^38,31^. This resemblance takes on added significance considering that experimentally induced activation of the PMd and the MAC trigger reaching-like movements of the arm in both monkeys and mice ^39,10,3^. Given that reach-to-grasp movements in rodents are considered innate and evolutionarily conserved, it is expected that the neural principles governing this behavior are shared across species ^40,8,41,42,43^. Thus, the converging evidence points to a possible degree of functional homology between the PMd of primates and the MAC of rodents.

In addition, optogenetic inactivation of the MAC, but not the primary motor cortex, impairs directional reaching in mice ^8^, supporting a potential direct role in the control of movements. At first glance, layer 2/3 neurons of the MAC could contribute to this function. For instance, target-related and reach-related neurons could encode movement goals towards the reaching target during the preparatory and executions phases of the task. However, this interpretation does not account for neurons active during drinking and intertrial periods, whose activity is equally tuned to the reaching target location. A way of reconciling these findings could be that layer 2/3 neurons perform a variety of independent functions, such as motor preparation, outcome processing and working memory ^33,34,44^.

Alternatively, we propose a more cohesive interpretation in which layer 2/3 neurons in motor areas perform a unified function where their activity is combinatorially determined by behaviorally relevant dimensions. In the case of the water reaching task, these dimensions include the position of the reaching target, the behavioral phase and the effector executing the task. Therefore, the population of layer 2/3 neurons in the MAC integrate real-time internal (somatosensory and motor) with external (spatial) information providing a cellular basis for high order representations of the animal’s actions^45^. In principle, this representational interpretation might seem at odds with the current views of cortical functioning ^46,47,48,35^. However, most models of cortical motor control emphasize the importance of feedback in guiding behavior. By continuously integrating sensory and motor feedback, the brain estimates the current state of the body within the environment, thereby enabling real-time adjustments to achieve the desired outcomes ^13,18,19,420,21,22^. While state estimation for reaching movements has been proposed to take place in the cerebellum and parietal cortex ^23,24^, the motor and premotor cortices are believed to use these signals to refine ongoing motor commands ^22^. In our experiments, the temporally low-pass filtered responses of GCaMP6s imaging limit our ability to investigate state estimation during rapid arm movements. Nevertheless, the conspicuous signals observed in layer 2/3 of the MAC highlight the possibility of an overarching state estimation mechanism operating at the task’s timescale. Such estimations would provide layer 5b, the output of the MAC, with contextual information regarding the state of the world (e.g. “the target is located on the left”) and of the animal (e.g. “mouse is drinking with the left hand”), critical for establishing upcoming actions ^16,49^. Therefore the activity of layer 2/3 could indirectly influence the generation of motor commands by conditioning the cortical activity in layer 5b to produce the appropriate output^50,25,14^ before reaching motor centers in the midbrain and spinal cord ^6,7^.

Finally, we suggest that a corollary function for these high-order representations could be to provide the animal with a cognitive map allowing for conscious perception of the executed actions, supporting a general role of layer 2/3 in cognition ^51,52^.

## SUPPLEMENTARY MATERIAL

**Supplementary Figure 1:**
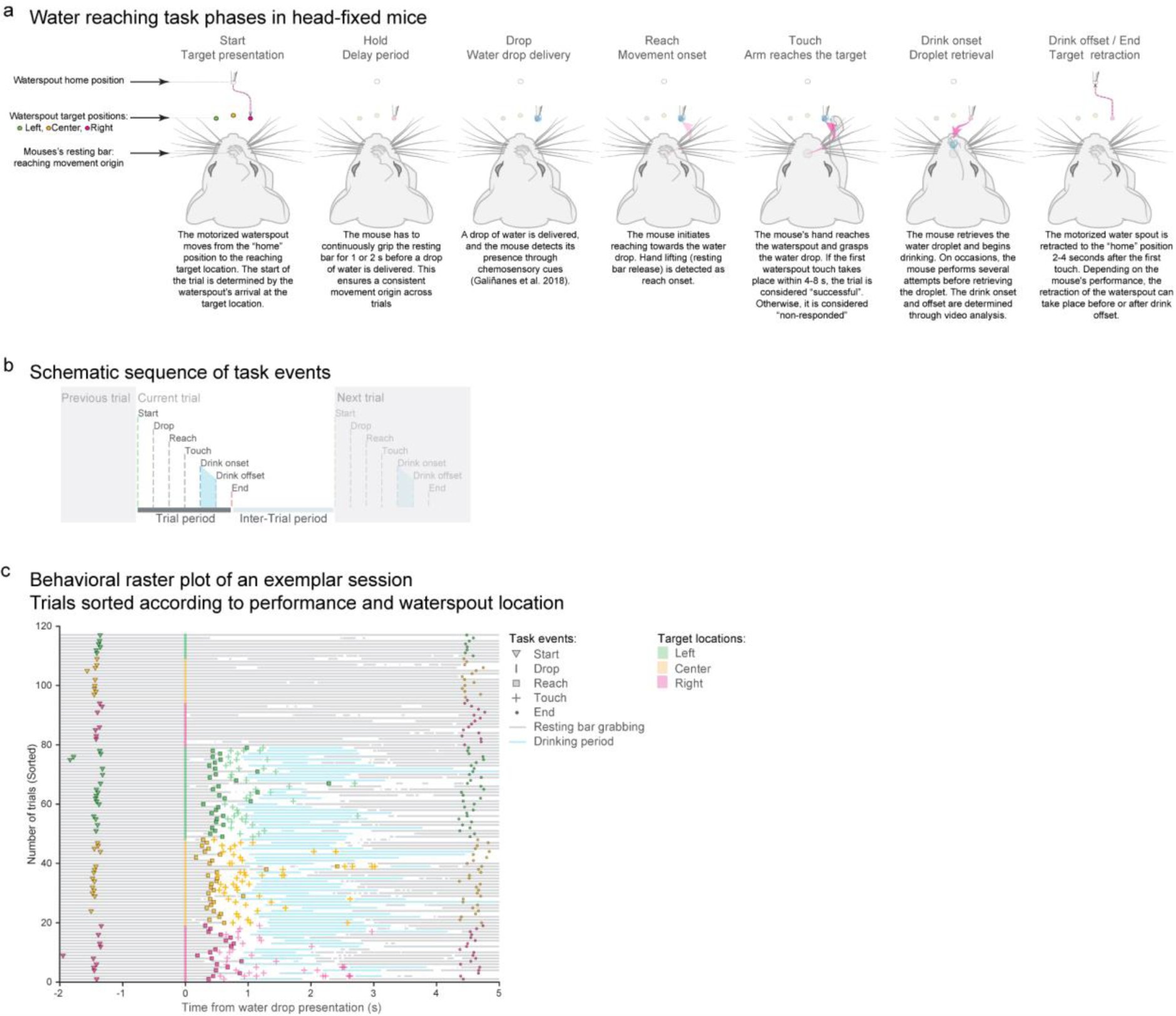
Water reaching task structure. **a** Schematic representation of the water reaching task for head-fixed mice during a successful trial. A new trial begins when the waterspout arrives in the reaching target location (target presentation). A delay period of at least 1 second is imposed before water drop delivery. In the dark, mice detect the location of the waterspout and the water drop using chemosensory cues ^8^. Following water delivery, mice perform directional reaching movements to retrieve and drink the water. The trial ends when the waterspout is retracted to the home position, marking the start of the inter-trial period. **b** Schematic representation of the task event timings, acquired either through the behavioral control system (drop, reach, touch) or via offline video analysis (start, drink onset, drink offset, and end). **c** Example raster plot depicting the behavior of a mouse during one session of the three-target water reaching task. Trials are aligned to the time of water drop presentation and sorted according to the outcome of the trial (successful or non-responded) and the trial type (target location; in this example target locations were Left, Center and Right). Different task events are coded with specific symbols. Horizontal gray lines indicate that the mouse’s hand was in contact with the resting bar. Triangles indicate the time of waterspout arrival and the beginning of a trial. Squares indicate the time of reach onset. Plus markers indicate waterspout touches. Horizontal blue lines indicate the drinking period. Dots indicate the end of the trial period and waterspout retraction. Note that on some occasions, mice perform multiple reach-to-drink movements to fully retrieve the water droplet.

**Supplementary Figure 2:**
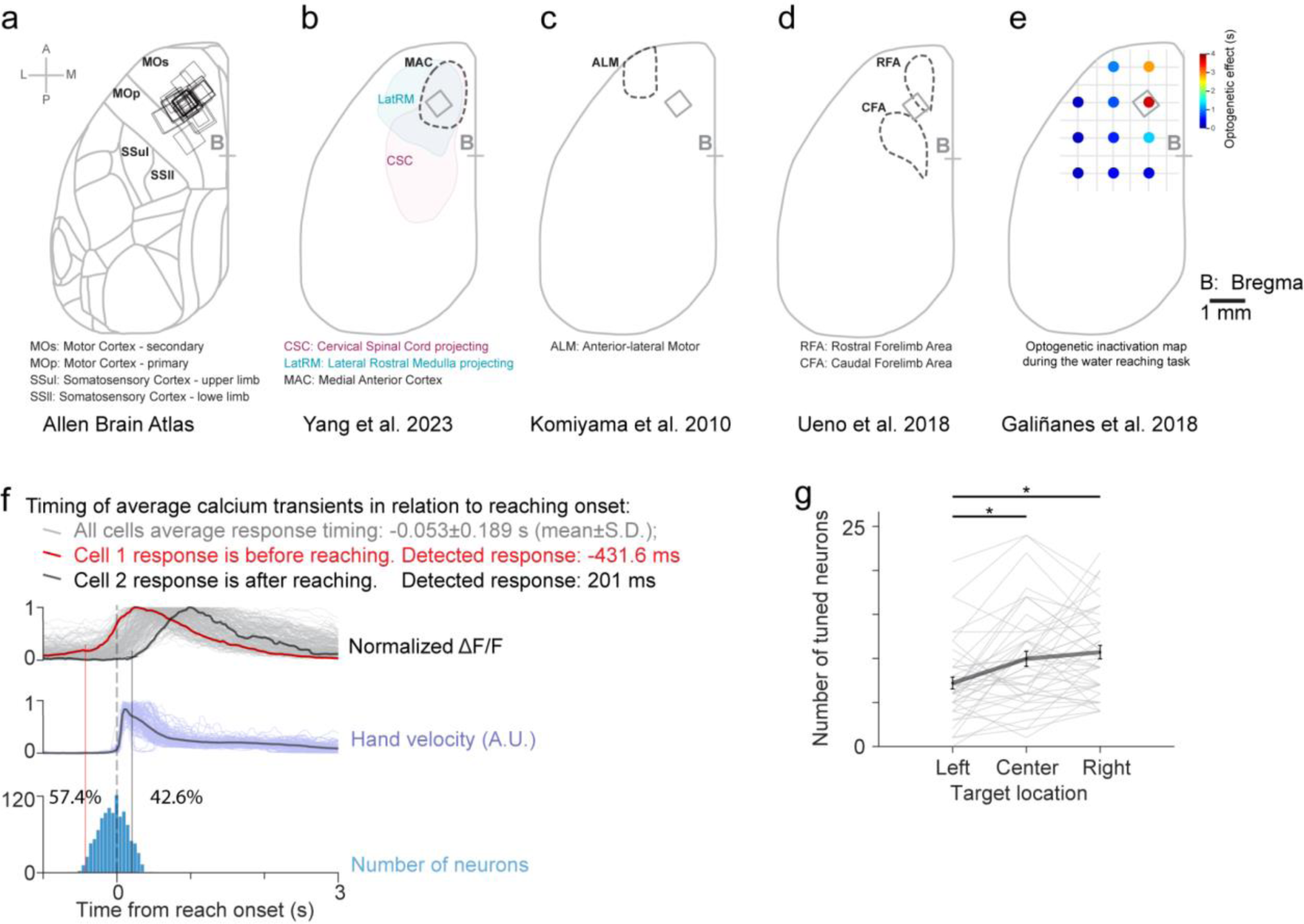
Two-Photon imaging of cortical neurons within the Medial Anterior Cortex (MAC) during the water reaching task. **a-e** Schematic representation of the dorsal aspect of the mouse cortex in relation to imaging FOVs; A: anterior, P: posterior, M: medial, L: lateral, B: bregma. **a**: Displays the main cortical areas as defined by the Allen Brain Atlas along with the fields of view (FOV) of the two-photon imaging sessions reported in this manuscript. Note that the FOVs are predominantly situated within the secondary motor cortex (MOs). **b** The medial anterior cortex (MAC) is determined by the cortical areas that project simultaneously to the cervical spinal cord and the lateral rostral medulla. The MAC region has been suggested to contain the cortical circuits involved in reaching ^6^. **c** The anterior lateral motor area has been involved in motor control of directional licking ^53^. **d** Caudal and Rostral forelimb areas are determined by the corticospinal projections to the cervical segments of the spinal cord ^54^. **e** Optogenetic inactivation map during the water reaching task showing that the maximal behavioral effect is obtained by inactivation of cortical spots within the MAC region ^8^. Gray square on the map corresponds to the representative FOV shown in the main Figure 1. **f** Reach-related neurons were detected as those neurons that responded within 500 ms prior to reaching onset and before touching the waterspout. Reach onset and waterspout touch were electronically detected (see methods). Top panel: The gray traces show the trial-averaged activity of reach-related neurons aligned to reach onset. Two example neurons are highlighted in red and black, which respectively respond before and after reach onset. Middle panel: trial-averaged hand velocity shows that the reach-related neuronal responses span from before movement (hand velocity equals zero) and during hand movement. Bottom panel: histogram of the response timing of reach-related neurons. **g** Number of reach related neurons responding to Left, Center and Right trials reveals a lateralization of the number of neurons in relation to the contralateral arm (left arm) executing the movement: there are more neurons that respond to reaching towards locations on the right hemispace of the animal. Gray lines individual sessions, black line population average, error bars: S.E.M. (N=44 sessions from 14 mice) *p<0.001

We also investigated whether the neuronal activity observed in the MAC was specifically tied to the arm used for reaching. To explore this, we compared the activity of reach-related neurons when mice reached using either their right or left hand ^38^. As shown in this supplementary figure, we found that a majority of reach-related neurons (46.5±5.2 %) were active only during right-hand reaches (i.e. contralateral to the recorded motor cortex) but were silent when the animal reached with the left hand. A smaller proportion of neurons (13.3±2.5%) were responsive during reaches made with either hand. Intriguingly, a sizable proportion of neurons (40.2±5.6 %) were active when the mouse reached with its left hand, which is ipsilateral to the recorded hemisphere. Importantly, all the neurons that were active when the mouse performed reaching movements with either of its hands, maintained their spatial tuning selectivity profile. This observation further supports the idea that these neurons encode external spatial locations. Taken together, our findings underscore that reach-related neuronal activity in the MAC is specifically linked to hand movements, effectively ruling out the possibility that the recorded activity is driven by whisker pad or nose movements.

**Supplementary Figure 3:**
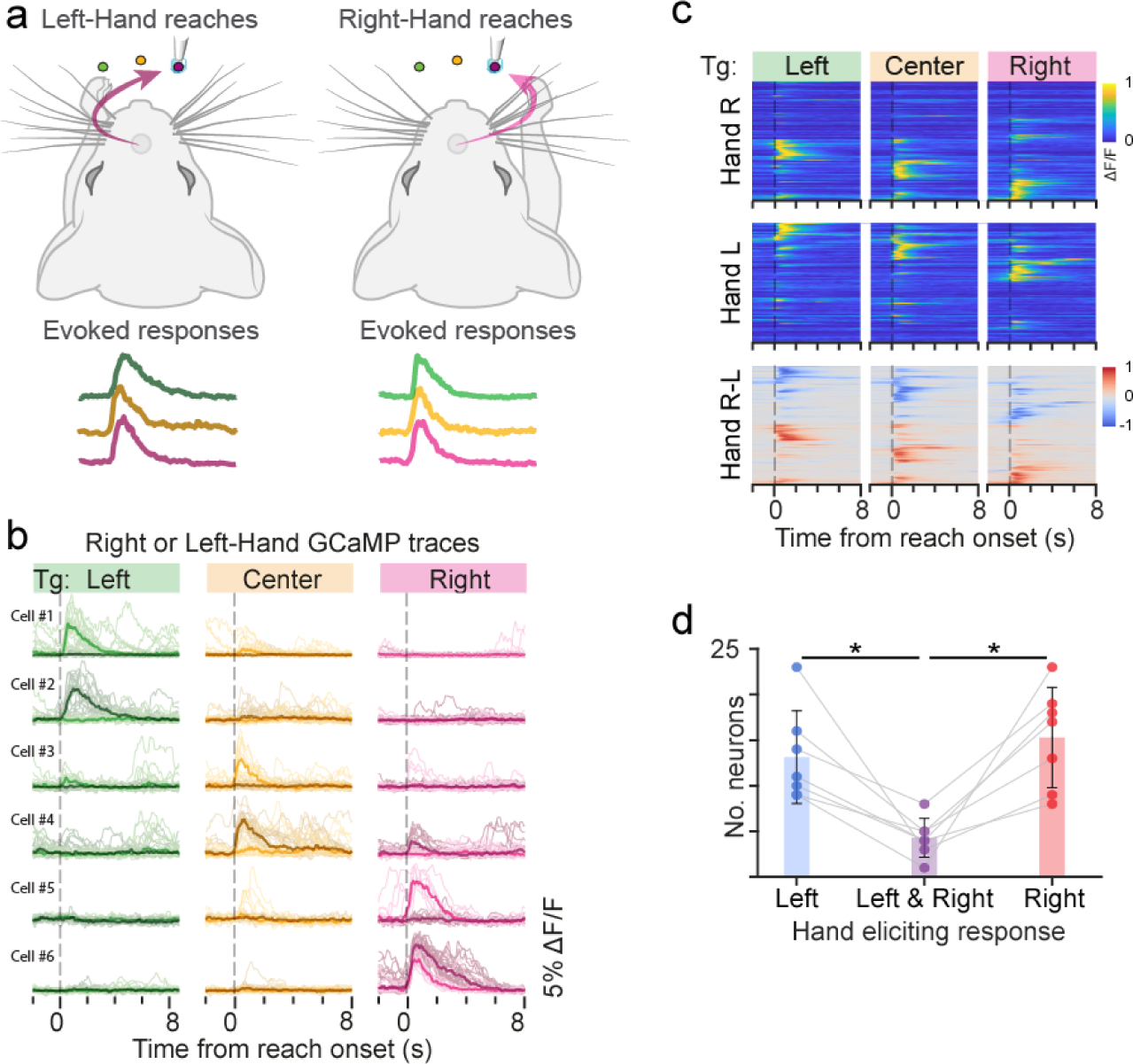
Cortical neuron activity is tuned to the arm executing the movement. **a** Schematic outline of the behavioral strategy: head-fixed mice reached either to the Left, Center, or Right target locations using either the right hand or the left hand. Blocks of 30-50 trials were alternated between Right-Hand and Left-Hand reaching during the same two-photon imaging session to compare the activity of the same neurons under both conditions. Two-Photon imaging recordings were always performed on the motor cortex of the left hemisphere. Calcium traces for Right-Hand reaches are depicted in light color shades while those of Left-Hand reaches are depicted in darker shades. **b** Similar to Figure 1g. Representative PECTs of six example neurons. Neuron activity during Left-Hand and Right-Hand reaching trials are overlaid for comparison. **c** The normalized PECTs of all reach-related neurons are divided into two panels based on the reaching hand used: “Hand R” for Right-Hand reaching trials, “Hand L” for Left-Hand reaching trials, and “Hand R-L” for the subtracted data between the two. **d** Number of active neurons during movements made with the Left hand, Right hand, or both. Bars: mean; error bars: S.E.M. * p<0.05

**Supplementary Figure 4:**
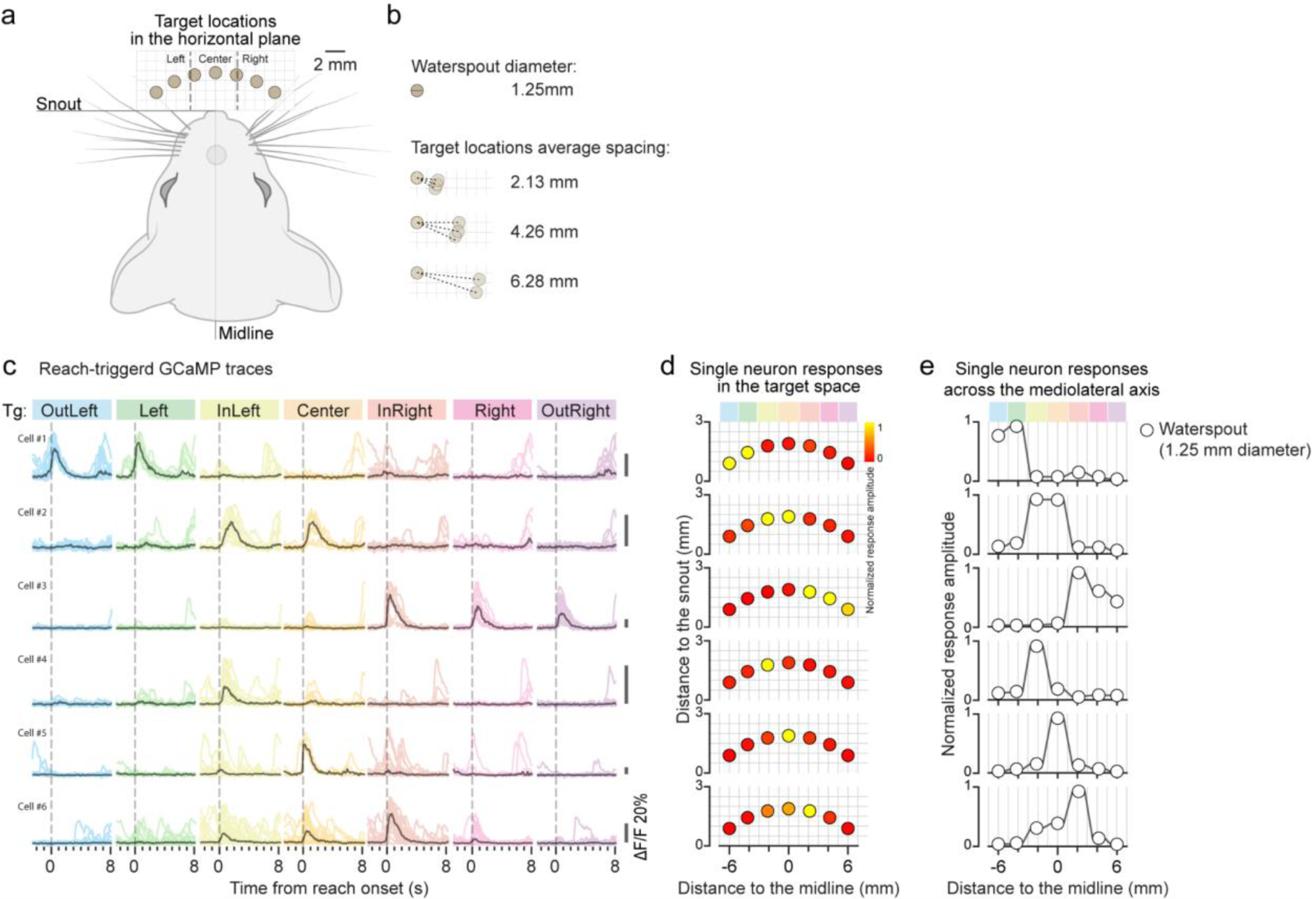
Sharp response fields boundaries at the single neuron level during the seven-target reaching task **a** Representation of the seven-target locations in the horizontal plane in relation to the mouse midline, tip of the snout. Waterspout diameter is represented to scale. Dashed lines represent the boundaries between Left, Center and Right response fields obtained in Figure 2g. **b** To-scale representation of the waterspout diameter (1.25 mm) and its relation to the spacing between targets locations. Adjacent target locations have an average separation of 2.13 mm, those once-removed are separated by 4.26 mm, and twice-removed by 6.39 mm. **c** Six example reach-related neurons displaying a spatial tuning profile with sharp boundaries. **d** and **e** The response amplitude of the neurons presented in (c) is plotted against their spatial relation to the target space (d) or across the mediolateral axis (e).

**Supplementary Figure 5:**
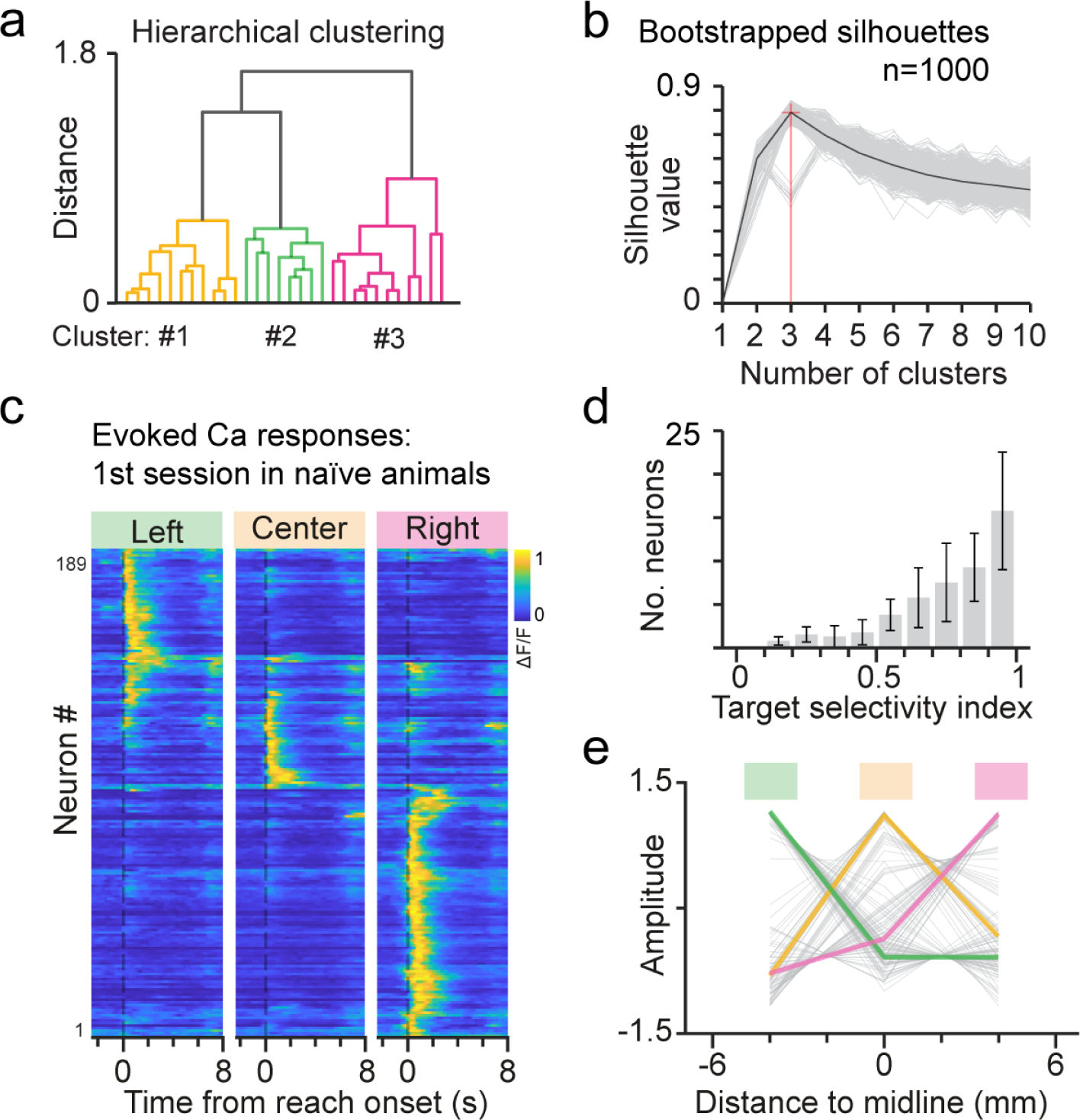
The target space division into three main response fields is an intrinsic characteristic of the MAC circuits. **a** Single-session dendrogram representing the hierarchical clustering of reach-related neurons based on the cross-correlation of their spontaneous activity (n=30 neurons simultaneously recorded in the session presented in Figure 2b). Each cluster was associated with reaching targets located towards the Left, Center or Right. **b** Unsupervised determination of the optimal number of clusters using the Silhouette scores. The Silhouette method assesses cluster quality by comparing an object’s similarity within its cluster to its similarity with neighboring clusters; a value close to 1 indicates good clustering, while a value near 0 suggests overlapping clusters. The seven-target response profiles (Figure 2g) were bootstrapped and clustered using 2 to 10 k-means clusters to compute the Silhouette scores. The analysis revealed an optimal number of 3 clusters (red line) with a 95% confidence interval. **c** Two-photon recordings in naïve mice during the first three-target reaching session revealed that the spatial organization into Left, Center and Right spaces is independent of training. After head-fixation habituation, naïve mice were exposed to the three-target water reaching task for the first time. The activity of MAC neurons was simultaneously recorded. Similar to expert mice, reach-related neurons of naïve mice displayed a strong target location selectivity for Left, Center and Right locations confirming that the reaching target space division is independent of training. Data shows the normalized average activity of all reach-related neurons acquired during the first three-target water reaching session (189 neurons, 4 mice, 4 FOVs). **d** Target selectivity index for the same neurons shown in (c). Bar:mean, errorbar: S.E.M. **e** Z-scored response amplitude profiles of individual neurons shown in (c, n=189, gray lines) across the mediolateral axis or the mouse. Colored lines: Average response profile for each of the k-means clusters color coded according to their spatial preference.

**Supplementary Figure 6:**
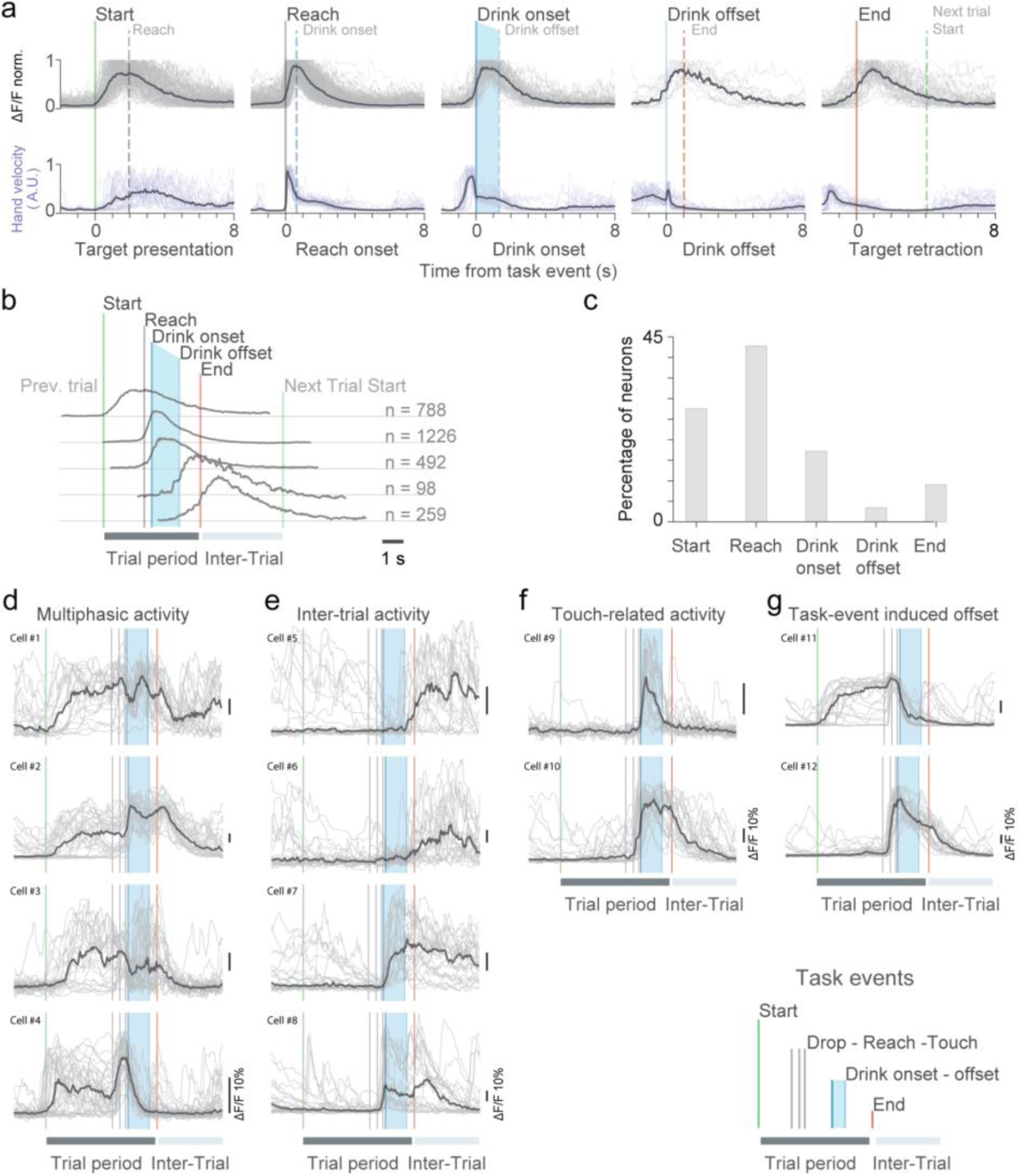
Task-related neurons active during different task events. **a** Top panels: PECTs (normalized fluorescence) aligned to relevant task and behavioral events ranging from the start to the end of the trial indicate that a substantial number of neurons in the MAC are active not only during reaching movements, but also during other specific events relevant the water reaching task. Thin gray traces correspond to the trial-averaged changes in fluorescence of individual neurons (20% of the full data set for visual clarity). Thick black traces are the population average (20% of the full data set). Solid vertical lines correspond to the time of the alignment event: target presentation, reach onset, drink onset and offset and target retraction. Dashed lines correspond to the average timing of the next behavioral event. Bottom panels: hand velocity (arbitrary units, normalized) in the same time scale of the calcium traces show the different behavioral state of the animal. **b** Average activity of neurons responding to different phases of the task (a) pseudo-aligned to the average timing of the behavioral events (vertical solid lines) as timepoint references. “n” indicates the number of recorded neurons in the full dataset. **c** Distribution of the recorded neurons according to their phase preference. **d-g** Time stretched calcium traces of 12 different neurons. Gray traces correspond to individual trials and black traces are their average. Vertical solid lines correspond to the task events used for time stretching: target presentation (green), drop delivery followed by reach onset and waterspout touch (gray), and target retraction (red). Blue shade corresponds to the average drinking period. **d** Example neurons that were active throughout the trial displaying sustained activity from target presentation up until retraction (Cell #1-#3). Some of the neurons show additional increases of activity at multiple phases of the task such as water drink onset (Cell #2) and drop delivery (Cell #4) indicating multiphasic responses. **e** Example neurons displaying sustained activity during the intertrial interval and in response to target retraction. **f** Example of touch-related neurons. Cell# 9 is considered a purely touch-related neuron. On the other hand, Cell #10 initially responds at reach onset but shows an additional increase of activity upon waterspout touch, which persists during drinking until waterspout retraction. **g** Example neurons with sustained activity that display a sharp fluorescence decrease (“offset”) upon specific task events suggesting a silencing mechanism. Cell #11: offset induced by waterspout touch. Cell# 12: offset induced by waterspout retraction.

**Supplementary Figure 7:**
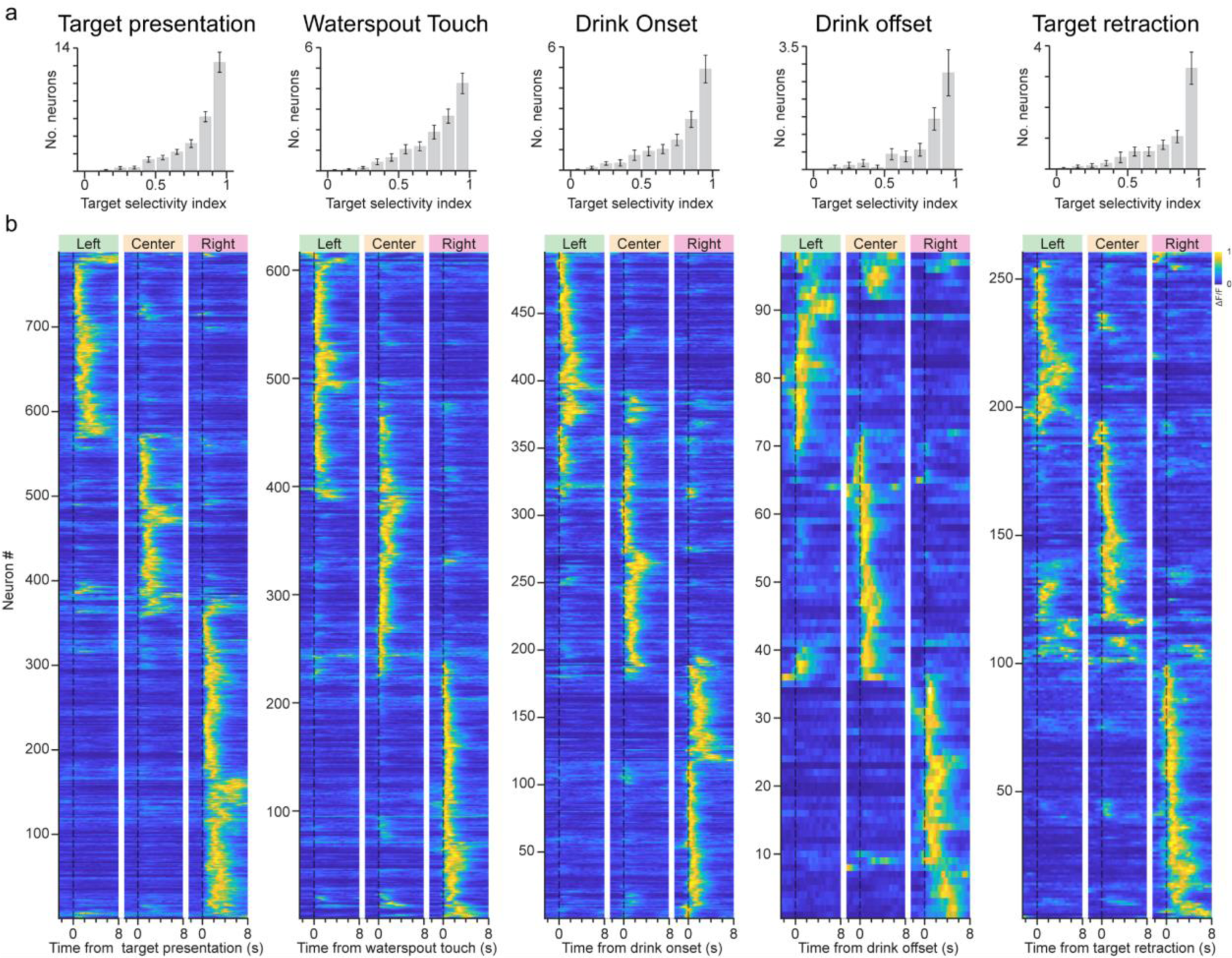
Neurons tuned to different task events are simultaneously tuned to the reaching target location. **a** Target selectivity index for recorded neurons tuned to different task phases: target presentation, waterspout touch, drink onset, drink offset and target retraction indicates that all task-related neurons display a similar selectivity for the waterspout location as reach-related neurons (Figure 1d). **b** Normalized average activity of all task-related neurons in response to reaching target locations on the Left, Center and Right. The data shows that task-related neurons conjointly encode the ongoing phase of the task (from target presentation to target retraction) with the location of the waterspout. Remarkably, neurons tuned to the target retraction event tended to display sustained activity during the intertrial period, when the waterspout was out of reach (supplementary figure 6) suggesting a continuous abstract representation of the reaching target space (i.e.: independent of sensory stimulation).

**Supplementary Figure 8:**
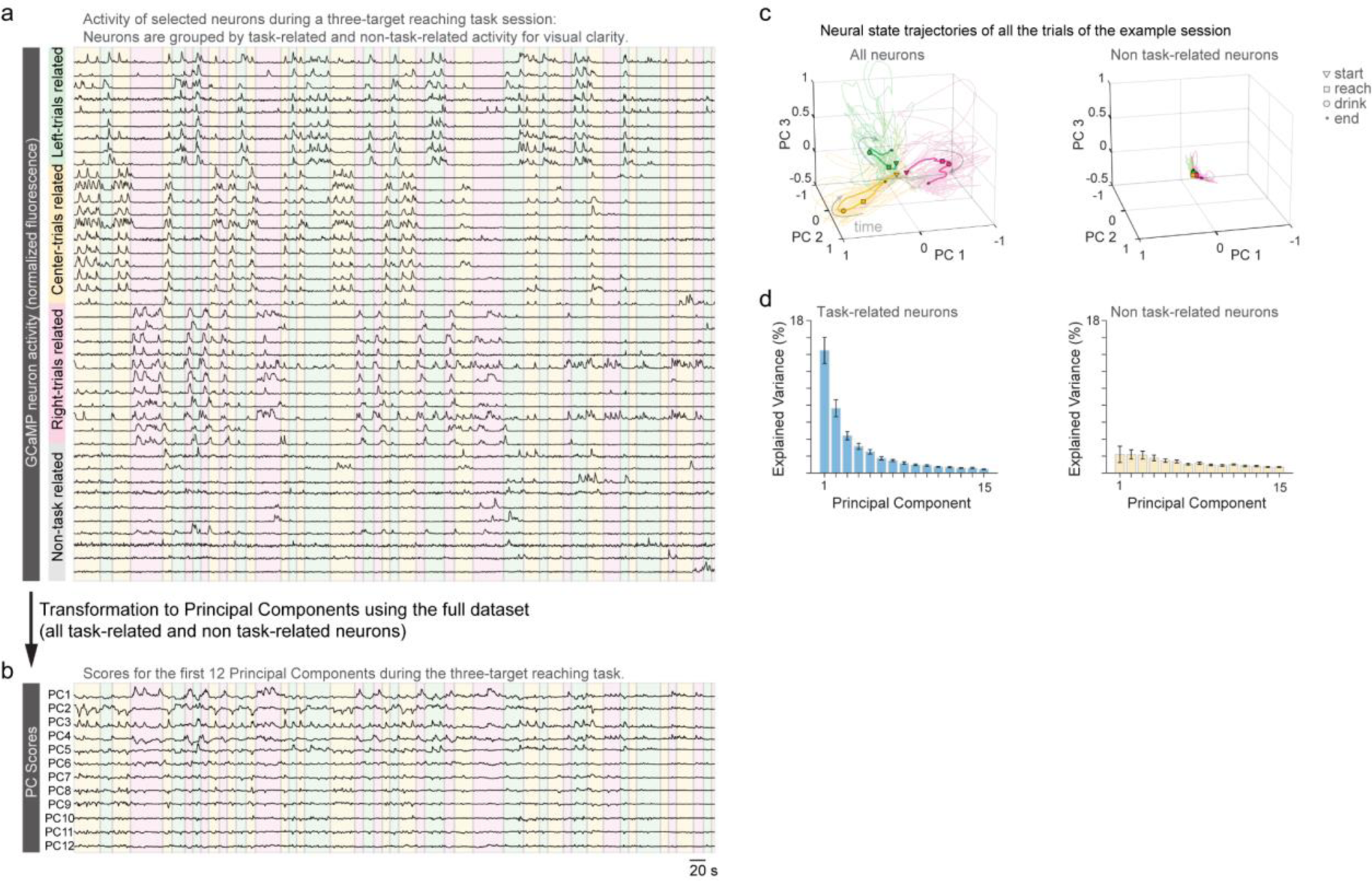
Principal components transformation of GCaMP fluorescence traces. **a** Fluorescence changes of a subset of task-related and non-task related neurons recorded simultaneously during a three-target reaching session. Individual neurons are sorted according to their target location tuning (Left, Center and Right), as shown in Figure 3a. Non-task related neurons typically displayed sparser activity than task-related ones. **b** The traces of all recorded neurons were transformed to principal components to obtain a dimensionally reduced representation of neuronal activity. The first 12 PCs are shown. **c** 3D representation of neural state trajectories of the example session shown in (a) across different time-stretched trials of the first 3 PCs. Thin lines correspond to individual trials and thick lines to their average. Color shades represent the waterspout location during the corresponding trials. Gray arrows indicate time progression specific task events (start, reach, drink onset and end) are displayed with markers on the average neural trajectories. Left panel: PC scores obtained from all recorded neurons (b) shows that the neural state trajectories diverge upon target presentation (start) following near orthogonal trajectories according to the waterspout location throughout the full trial length; Right panel: the activity of non-task related neurons was projected to the same PC space. Neural state trajectories of non task-related neurons appear stagnant, remaining around the coordinate origin, revealing that the neuronal state trajectories are not contributed by non-task related neurons. **d** Explained variance for each of first fifteen PCs for task-related neurons and non task-related neurons further confirms that non-task related neurons do not carry significant information during the execution of the water reaching task (10 mice, 27 FOVs). Bars: mean, error bars: S.E.M.

**Supplementary Figure 9:**
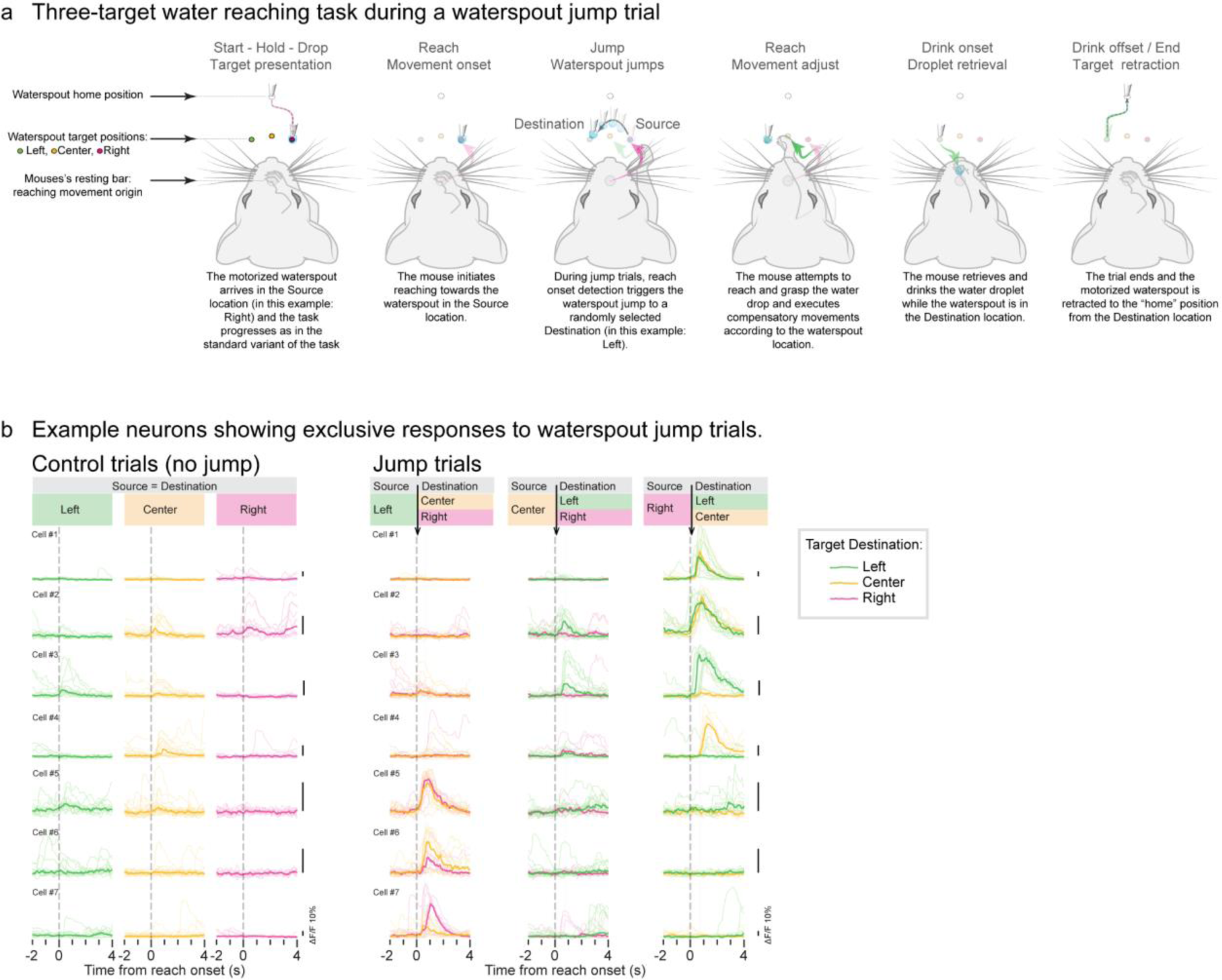
Schematic representation of the waterspout jump task variant. **a** Schematic representation of the waterspout jump variant of the three-target reaching task for head-fixed mice during a “jump” trial. The initial phases of the trials (from start to drop) were equivalent to the standard variant of the task. During “jump” trials, the waterspout was rapidly (25 mm/s) relocated from the initial target position (“Source”) towards another target location (“Destination”) which was randomly chosen between remaining targets (e.g.: if the source location was at the Right, the destination could be Left or Center). Waterspout jumps took place after reach onset was detected. Upon waterspout jump, the mice spontaneously (no training required) performed adjusting movements to reach the drop of water in the current location. On some occasions, mice performed multiple reaching attempts before retrieving the drop of water. In some trials, mice managed to retrieve the drop of water when the waterspout was still at the “source” location. **b** A subset of neurons in the MAC are not active during control trials (left panel) but respond in jump trials (right panel). PECTs on the left and right panel correspond to the same neurons. The waterspout Source/Destination locations are indicated by the Left, Center and Right columns. The color of calcium traces (thin lines individual trials, thick lines median average) also indicate the waterspout Destination location (green jumps to Left, yellow jumps to Center and pink jumps to Right). Black arrows indicate the average time of waterspout jump. PECTs are aligned to reach onset (dashed line). Cell #1 and #2 respond to jumps from Right to either Left or Center locations. Cell # 3 responds to jumps from Right and Center to Left. Cell #4 responds to jumps from Right to Center. Cell #5 and #6 respond to jumps from Left to either Center or Right. Cell #7 responds to jumps from Left to Right.

## METHODS

### Animals

All experiments were approved by the Animal Care Committee of the University of Geneva and by the “Direction générale de la santé” of the Canton of Geneva. Mice were held under a controlled 12/12 h light/dark cycle (7:00 h lights on, 19:00 h lights off) with ad libitum access to food and water until the start of behavioral experiments. Mice were group housed and provided with nesting material (shredded paper or cotton), cardboard houses and wooden gnawing blocks to improve welfare. Animal wellbeing before and during experimental manipulations was ensured by following recommended laboratory animal practices ^55^. Experiments were performed on adult (12-30 week old) mice of both sexes: 13 female and 2 male double transgenic mice Camk2a-GCaMP6s (generated by crossing heterozygous B6.Cg-Tg(Camk2a-tTA)1Mmay/DboJ and B6;DBA-Tg(tetO-GCaMP6s)2Niell/J mice). Camk2a-GCaMP6s express the neural activity indicator GCaMP6s ^56^ in excitatory neurons of the cortex ^57^. Wild type mice were purchased from Charles River and transgenic mice were obtained from The Jackson Laboratory and bred in the animal facility of the University of Geneva.

### Surgeries

Head-bar and cranial window surgical procedures for two-photon imaging were performed as previously described ^58^ and following recommended refinement procedures ^55^. All surgical interventions were performed under general anesthesia and local (“wound level”) aseptic conditions. A balanced anesthesia regimen was administered for pain and inflammation management before, during and after surgical procedures. Prior to general anesthesia, mice received a subcutaneous [s.c] dose of 0.1 mg/kg buprenorphine. Twenty to forty minutes later, general anesthesia was induced with gaseous isoflurane (5% for 1 to 2 minutes). After nociceptive reflex responses (toe and tail pinch) disappeared, the mouse was secured onto a custom-made stereotaxic frame equipped with an anesthesia mask (isoflurane maintenance at 1.5%) and received analgesic and anti-inflammatory drugs (2.5 mg/kg intramuscular dexamethasone, and 5 µL s.c. of 1% lidocaine under the scalp) to reduce brain swelling and pain. The mouse’s body temperature was maintained at 37°C with a heating plate. To prevent corneal damage the eyelids were closed with petroleum jelly. Five minutes after lidocaine injection, the scalp was cleaned (ethanol 70%), disinfected (chlorhexidine 0.5%) and excised. The exposed periosteum was removed using sterile cotton swabs and corneal scissors. Fiduciary marks on the skull were made at the level of bregma and the midline to provide reliable reference points for future imaging experiments. The marks were performed by making small incisions with a scalpel and permanently staining them with a waterproof marker. An oval shape craniotomy of approximately 2.3 by 4.2 mm was delineated around the medial anterior cortex (MAC) region of the secondary motor cortex of the left hemisphere to image the activity of reach-related cortical circuits ^8,6,3^. The surface of the skull was gently scraped to increase adherence and covered with a thin layer of cyanoacrylate glue (ergo 5011 or Loctite 401). The skull around the craniotomy area was thinned with a dentist drill equipped with a cylindrical bur. During this procedure, the skull was bathed with sterile saline at room temperature to soften the bone, reduce bleeding and prevent excessive heating of the brain. Once the skull was thinned down to approximately 100 μm, the craniotomy was performed using a scalpel. The cortical surface was rinsed with sterile saline and bone debris were removed before replacing the excised bone with a glass cranial window. The cranial window, matching the shape of the craniotomy, was made out of two hand-cut coverslips (150 μm thick) using a diamond scribe (Fiber Instruments, FO90C) and stacked together with UV glue (Norland 61). The cranial window was gently positioned on top of the brain and secured against the bone with cyanoacrylate glue. A custom made titanium head-bar was positioned on top of the interparietal bone and all the implants were covered with clear acrylic resin (Ortho-Jet resin powder and liquid, Lang). At the end of the surgery and for the 2 to 5 following postoperative days, mice received a daily dose of 5 mg/kg carprofen s.c. for prolonged analgesic and antiinflammatory effects. Mice were typically housed in groups of 2 to 4 and left to recover from surgery for at least 5 days before behavioral manipulations.

### Behavior: naturalistic water reaching task

Mice performed a directional water reaching task adapted from a previously described task ^8^. Briefly, after recovering from surgery, mice were water deprived and entered the handling period. Daily handling was performed following published recommendations ^55^ and consisted of gradual habituation to the experimental area and head-fixation setup. The head-fixation setup consisted of a 3D printed mouse enclosure and a pair of custom-machined steel headbar clamps. The clamps were held at an angle of 8° in the sagittal plane and elevated 40 mm from ground level allowing the mice to remain in a standing position during head-fixed behavior. The 3D printed enclosure was designed to minimize animals’ discomfort, promote natural body postures and allow for a controlled manipulation of the behavioral output. Habituation to head-fixation was achieved progressively by increasing head-fixation duration at a rate of ∼5 minutes per day, while simultaneously providing the mice with tap water (up to 1 ml) to counteract head-fixation discomfort. During this period, mice were encouraged to grab a resting bar located close to the body midline with their right hand while a 3D printed paw blocker limited the use of their left hand. During this period, water was manually provided through a syringe coupled to a blunt needle, which the mice licked from. Unless stated otherwise, mice were typically induced to reach and grasp water droplets by placing the needle away from their mouth.

After the habituation period (5-7 days), mice started the automatized water reaching task in the dark. In this task (Supplementary figure 1), inspired by the primate center-out reaching task, head-fixed mice performed directional reaching movements from a starting point (“Origin”) towards a reaching target located around their snout. The reaching origin was determined by the resting bar position and the reaching target location by a mobile waterspout which provided water droplets. At the beginning of each trial, the waterspout was at a distant “home” location (5 cm away from the snout aligned within the midline of the animal) and moved rapidly (25 mm/s) to the “target” location. Target locations were pseudo-randomly selected from a predetermined set of coordinates within the horizontal plane ∼5 mm below the tip of the snout. In order to trigger the droplet presentation, mice were required to continuously touch the resting bar for a delay period of 1 or 2 seconds. Releasing the resting bar before that period, reset the timer. If the mice didn’t touch the resting bar within a window of 6-10 seconds, the trial ended. In the dark, mice detect the presence of the waterspout and water droplet using chemosensory cues and typically reach for, grasp, retrieve, and drink the water within 2 seconds ^8^. Therefore, the waterspout was left in the target position for 2-4 seconds after the initial touch before the trial ended. However, if mice failed to touch the waterspout within 4-8 seconds after droplet presentation, the droplet was suctioned away and the trial ended. At the end of each trial, the waterspout returned to the home position and a new trial began. Therefore, in addition to reaching, the water reaching involves multiple well defined phases determined by specific task events with which the animal interacts with: target presentation followed by a delay period (Start), movement response (Reach), water consumption (Drink onset and offset) and target retraction followed by an intertrial period (End).

We would like to highlight that, in contrast to the majority of primate reaching tasks where movements are typically performed through a manipulandum or towards virtual reaching targets ^59^, the water reaching task is naturalistic, providing greater affordances as it allows head-fixed mice to interact with a graspable physical object that enters and exits their peripersonal space mimicking the dynamics of real-world experiences. The animal’s output is a cohesive reach-to-grasp-to-mouth sequence of movements. These aspects add an ethological value to the task, which, except for the head fixation, resembles the types of movements and interactions that the mice would likely encounter in their natural environment. Therefore, the neuronal correlates of the task probably reflect innate neuronal circuits for behavioral control.

To probe for different coding aspects of cortical neuron activity, we introduced a series of variants to the standard task: (1) Alternative reaching origins, (2) Arm switch, (3) Seven-target reaching, and (4) Waterspout jump.

#### Alternative Reaching Origins

To determine whether cortical neurons encoded kinematic aspects of reaching movements, we analyzed the neuron activity during trials in which reaching movements were initiated from two alternative origins. This manipulation was achieved by displacing the resting bar using a linear stage (T-LSM100A, Zaber Technologies). The standard origin of reaching was typically aligned to the midline of the animal (Origin A) and the alternative origin (B) was 8 to 10 mm away in the mediolateral axis. During a given session, several blocks of 10-30 trials from each origin were probed to compare the activity of the same neurons under both experimental conditions.

#### Arm Switch

All mice used for two-photon imaging underwent cranial window surgery on their left hemisphere. In the standard task, mice were trained to use their right hand for reaching. This was achieved by placing a paw blocker in front of the left hand. To determine if neuronal activity was tuned to the hand executing the reaching movements, mice were also trained to reach using their left hand by switching the paw blocker’s position. Achieving comparable performance levels with Left-Hand reaching typically required 3 to 5 training sessions. During “arm switching” sessions, mice performed at least two alternate blocks of Left-Hand and Right-Hand reaching, with each block typically consisting of 30-50 trials.

#### Seven-target reaching

After training in the three-target reaching task, mice were introduced to a seven-target reaching task in order to investigate the fine-scale spatial tuning properties of cortical neurons. While the original targets in the standard task (Left, Center and Right) were spaced by 4.2 mm, the expanded version added four novel target locations (Outer-Left, Inner-Left, Inner-Right, and Outer-Right), thereby reducing the average target spacing to approximately 2.13 mm (Supplementary figure 4). Most mice adapted to this task variant did not require any specific training. However, because Outer-Left and Outer-Right locations were at the edge of the maximal reaching space ^8^, some mice did not perform as well to these target locations.

#### Waterspout Jump

To investigate how changes in the reaching space are processed by cortical neurons, we introduced a variant to the standard three-target reaching task. In this variant, the waterspout’s location was unpredictably altered during an ongoing trial. For control trials (“no-jump”), the waterspout’s behavior was the same as those of the standard task. Conversely, in “jump” trials, the waterspout arrived in a specified location (“Source”) and, shortly after reach onset, it was relocated (“jumped”) to one of the other two target locations (“Destination”; supplementary figure 9). The waterspout relocation event occurred in two-thirds of the trials chosen at random. In order to investigate all 9 possible Source-Destination combinations for the three-target reaching task, mice had to perform between 120 and 200 trials. It is important to note that this variant of the task was performed spontaneously by the mice without any specific training and mice were allowed to perform multiple attempts before water drop retrieval.

### Behavioral control system

The behavioral task was controlled using a real-time linux state machine through a MATLAB interphase (https://brodylabwiki.princeton.edu/bcontrol). The real-time system controlled various actuators such as solenoid valves (LHDA1231215H, The Lee Company), linear stages (T-LSM100B and T-LSM100A, Zaber Technologies) and LEDs. It also recorded touches of the resting bar and waterspout through transistor based sensors^60^. A custom made double lumen waterspout was made using a pair of nested 18G and 20G needles for water droplet delivery and suction. High speed usb cameras (MQ013RG-ON, Ximea) equipped with 5 mm lenses (Navitar) were used to video record the movements of the arm. 320 by 300 pixels images were acquired at 200 fps using Streampix 7.0 (Norpix) and arm trajectories were offline reconstructed using DeepLabCut^61^. Experiments were performed in the dark under infrared (IR) illumination (IR-LED ELJ-810-629, Roithner and IR-LED arraysM120, Kemo Electronic).

### Two-Photon, widefield fluorescence and image processing

Two-Photon imaging of layer 2/3 neurons was performed using a custom built two-photon microscope (MIMMS; https://openwiki.janelia.org/wiki/display/public/Home) controlled by Scanimage 5 (https://vidriotechnologies.com/). A 16x 0.8 NA objective (Nikon) with a pulsed laser excitation wavelength at 920 or 940 nm (Ultra II, tunable Ti:Sapphire laser, Coherent) and a resonant scanner system (Thorlabs) were used to image GCaMP6s fluorescence changes within fields of view (FOVs) ranging from 450 to 650 µm per side and captured at 29.38 frames per second on averge. Laser power was modulated by a pockel cell (350-80-LA-02, Conoptics) to a maximum power of 40 mW at the end of the objective. 512 by 512 pixel images were continuously acquired using a gated photomultiplier tube (H11706P-40 SEL, Hamamatsu) and digitally written in 16 bit format to disk in separate files triggered by the behavioral system using TTL pulses. For statistical purposes, imaging sessions typically lasted between 60 to 160 consecutive trials depending on the behavioral manipulations and up to 4 different FOVs were imaged during the same behavioral session. Except for the “naïve” mice experiment (Supplementary Figure 5e), two-photon imaging recording sessions started after mice were pre-trained to proficiently perform the water reaching task. A path for a CCD camera with the center of the field of view aligned to that of the two-photon microscope was integrated into the microscope setup. The CCD camera path was illuminated with green LED to obtain well contrasted reference images of the cortical blood vessel map across sessions and the fiduciary marks performed on the skull during surgery. CCD camera images were obtained using custom MATLAB code to precisely identify the location of bregma and the imaging coordinates. The imaging plane of the microscope was oriented parallel to the cranial window for each animal^58,62^. The imaging FOVs were located within the MAC region (∼1.5 mm anterior to bregma, ∼1.25 mm lateral to the midline) at different depths (ranging from 150 to 300 µm below the cortical surface). Images were processed with adapted scripts from the CaImAn toolbox for MATLAB^63^. Automatically detected regions of interest (ROIs) and extracted GCaMP fluorescence traces were curated and post processed with custom written MATLAB scripts. CaImAn parameters were empirically determined by iterations of extraction and visual curation of ROIs and calcium traces.

To obtain anatomo-functional reference maps, the cortical sensory representations of the right forelimb and hindlimb were mapped through widefield fluorescence imaging (Camk2a-GCaMP6s mice) and somatic vibratory stimulation. After the mice had fully recovered from the cranial window surgery, they were anesthetized with isoflurane and received a vibrotactile stimulation (1s at 100 Hz for 20 to 40 repetitions) on the forelimb or hindlimb while imaging fluorescence changes through the cranial window at 10 fps (256 by 332 pixel resolution, 12-bit Retiga EX camera, QImaging). The surface of the cranial window was illuminated with a blue LED light (470 nm, ROITNER) and the optic path was arranged with a set of FITC excitation and emission filters (NIKON). Stimulation and image acquisition were controlled with Ephus ^64^. Images were processed with custom MATLAB scripts by subtracting the baseline fluorescence of the pre-stimulation period (1 s average) to all the frames of each trial. Baseline-subtracted images were then averaged across trials and the peak fluorescence response within 1 s after the end of the stimulation was used to delineate primary somatosensory regions of the cortex.

### Calcium Data Analysis

#### Task-related neurons

To determine the presence of task-related neurons, we denoised the calcium traces with two consecutive moving average window filters (9-datapoint median followed by a 3-datapoint mean). We then screened the neuronal activity and performed perievent calcium traces (PECTs) by aligning the extracted fluorescence signal (ΔF/F) to a series of observable task events (“triggers”): waterspout presentation, drop delivery, reach onset, waterspout touch (first touch after drop delivery), drink onset, drink offset, and waterspout retraction. The timing of these task events was obtained from the behavioral control system or extracted from video analysis using DeepLabCut. We compared the normalized fluorescence of the extracted ROIs using a paired t-test based on the time-averaged fluorescence intensity from a window of 1 s before and after each triggering event. An ROI was identified as related to a specific phase of the water-reaching task if it displayed a significant response. This response was determined by a p-value below 0.01, an amplitude > 4%, an onset of calcium events within 300 to 1000 ms around the task event, and an increase in the ΔF/F slope before and after the event. Neurons were classified as reach-related only if their onset took place before the waterspout touch, thereby ruling out any touch-related influence. Using this method, we classified task-related neurons into the categories: start, reach, touch, drink, and end.

#### Spatial tuning, Selectivity Index and Clustering

To assess spatial tuning, task-related neurons were categorized based on the number of targets eliciting a significant response. Neurons were then grouped according to the number of targets eliciting response, with categories ranging from 1 to 3 or 1 to 7 depending on the number of reaching target locations. Inspired by the orientation selectivity index, we also computed an unbiased measurement of spatial tuning called the target selectivity index (TSI). The TSI quantifies the degree of response selectivity for each neuron using the formula: TSI = 1-(mean(non-preferred response)/preferred response), where ‘preferred’ refers to the highest response amplitude and ‘non-preferred’ includes all the others. A TSI of 1 implies that the neuron responded maximally to only one target location and had no response to all the others. Conversely, a TSI of 0 indicates equal response amplitude across all target locations. Similarly, to assess neuronal tuning based on different starting points of reaching movements, we devised an origin selectivity index.

To further describe the tuning patterns of task-related neurons, we applied clustering analyses, focusing on both the neuronal spontaneous activity and the response amplitude patterns. For the analysis of spontaneous activity, we generated a pairwise cross-correlation matrix from the neurons’ activity throughout the behavioral session. Hierarchical clustering was employed using the farthest distance method, with clusters delineated using the median distance as a cutoff.

Next, we established the response amplitude pattern of individual neurons by assessing the response amplitude elicited by each reaching target location. These amplitude profiles were standardized via z-score normalization, setting the stage for k-means clustering. The optimal number of clusters was determined using the elbow method (optimal number of clusters 3, not shown). To visually represent the clustered data, we transformed the full set of response patterns to its principal components (PCA) and plotted them in a three-dimensional space, utilizing the first three principal components. Each data point in this 3D projection was color-coded based on its k-means cluster designation and tuning bias. Finally, to ensure an unbiased determination of the optimal number of clusters, we implemented a bootstrap resampling strategy with 1000 resamples. Each resampled dataset underwent k-means clustering, examining a potential cluster range from 2 to 10. We next computed the Silhouette score for each bootstrap, and determined the highest Silhouette score as the optimal number of clusters ^65^. These values were used to establish a 95% confidence interval for the optimal number of clusters.

#### Trial alignment

Owing to behavioral variability, task events and durations of behavioral phases are inherently inconsistent, making straightforward alignment of full-length trials unfeasible. To simultaneously visualize neurons responsive to different task events, we employed a pseudo-alignment technique on the PECTs. For each neuron tuned to a particular event, we generated a corresponding PECT aligned to that event. We then determined the trial-averaged timings of the events that occurred before and after the primary event. Using the averaged task event times, we aligned traces across neurons. This technique allows the visualization of each neuron’s activity relative to its most closely tuned event, while preserving the temporal relationship between simultaneously recorded neurons. For quantitative comparisons of population activity across trials, sessions and mice, we implemented a time-stretching methodology. Each trial was segmented into discrete time series based on relevant task events: target presentation, water drop delivery, reach onset, waterspout touch, target retraction and next trial start. Subsequent to this division, each segment was linearly interpolated to adjust its length to match the standardized average duration of the associated behavioral phases across the whole population. This approach ensured trial uniformity in length across sessions and conditions, enabling inter-trial and across subjects comparisons.

#### Dimensionality reduction

For a comprehensive visualization and comparison of population activity, we employed the Principal Component Analysis (PCA; see supplementary figure 8). The full dataset of calcium traces from each session was transformed into its principal components. Before this transformation, fluorescence time series were denoised with a Gaussian filter, using a time window of 15-datapoints. After transforming the data, trials underwent time-stretching and were visualized in a three-dimensional space, to reveal the neural state trajectories throughout different task phases and behavioral conditions. To compare neural state trajectories across different sessions, we employed the Procrustes method to align and re-orient the first 15 PCs of the PCA results. Angles between trajectories of different conditions (i.e. target location) were calculated using the dot product of the normalized scores of the first 15 PCs.

To compare non-task related neurons and determine whether they convey significant information, the PC transformation was initially performed on the full data set of control trials and the dataset of non-task related neurons was further projected into the same space for comparison. Similarly, in the waterspout jump experiments, the PC transformation was initially performed on the control trials alone (i.e. “no-jump” trials) and the “jump” trials were projected into that space.

To determine the temporal evolution of neural state trajectories and compare them across conditions, we calculated the Euclidean distance between pairs of conditions using bootstrapped average trajectories derived from the first 9 PCs. This method illustrated the temporal evolution of the distance between different conditions (e.g., Right trials vs. Left trials or control trials vs. jump trials) as well as, through resampling, established the within-condition distances (e.g., Right trials vs. Right trials).

### Video analysis

Video recordings were processed using a DeepLabCut model trained on thousands of images to reliably reconstruct reaching trajectories, determine the drinking probability and the position of the resting bar and the waterspout across mice and sessions. The right hand of the mice was reconstructed using five “body parts”, with one marker for the wrist and one for each digit. Body parts extracted with a confidence value below 0.6 were excluded. Gaps in the data, resulting from missing body parts and spanning up to 10 consecutive data points, were interpolated using the ‘pchip’ method. Each body part was denoised with two consecutive filters: a 3-datapoint median filter followed by a 3-datapoint Gaussian filter. To represent the hand’s trajectories during reaching, the positions of all body parts associated with the hand were averaged in both X and Y dimensions. Velocity profiles for both the hand and the waterspout (expressed in pixel units) were determined using the gradient of their X and Y positions. The magnitude of the velocity was obtained by computing the sum of squared velocities in the X and Y directions. This velocity data was subsequently denoised using a 20-datapoint moving median filter. The velocity of the waterspout was used to determine the start (target presentation) and the end (target retraction) of each trial. The stereotyped velocity profile of reaching movements was used to align reaching trajectories across trials before computing average trajectories. The angle of reaching trajectories (θ) was determined by fitting a linear regression to the initial part of the reaching movement, from the onset to the peak velocity. The angle was computed against the vertical axis, where positive angles denote a rightward movement deviation. The drinking probability was extracted from the kinematic signature of the reconstructed body parts. The factors used were the orientation of the hand, the opening of the fingers, the vertical distance to the resting bar and the velocity of the hand.

### Statistics and data analysis

No statistical methods were used to predetermine sample size and all experimental animals were included in the analysis. Data were analyzed using custom-written routines, built-in functions, and published open-source toolkits in MATLAB and Python. Unless stated otherwise, average values of the neuronal population are represented as mean ± S.E.M. All statistical tests were performed using MATLAB. The Friedman test was used for comparison of repeated measures, followed by Wilcoxon signed-rank tests for pairwise comparisons. Categorical distributions were analyzed using a Chi-Square test. Adjustments for multiple comparisons were made using the Bonferroni correction to a significance level of p = 0.05.

### Data availability

Analyzed data and custom MATLAB routines are available from the corresponding author upon reasonable request.

## Acknowledgements

We thank M. Prsa, C. Schafer, and K. S. Lee for advice and comments on the manuscript. This work was supported by the Swiss National Science Foundation (310030_184829), the European Research Council (OPTOMOT), and the International Foundation for Research in Paraplegia.

## Author contributions

G.L.G. and D.H. conceptualized the study. G.L.G. designed, ran the experiments, wrote the code and analyzed data. G.L.G. and D.H. wrote the manuscript.

