## Supplementary Tables for "Continuous estimation of reaching space in superficial layers of the motor cortex"

**Extended Data Table 1: number of experiments**

| **Figure** | **Experiment** | **N animals** | **N sessions** | **N units** |
| --- | --- | --- | --- | --- |
| 1c | Spatial tuning of reach-related neurons | 14 | 44 | 1226 |
| 1d top | Target selectivity index of reach-related neurons | 14 | 44 | 1226 |
| 1d bottom | Targets eliciting response | 14 | 44 | 1226 |
| 1f | Reach trajectory angles | 6 | 6 | N/A |
| 1g | Reaching-origin tuning of reach-related neurons | 6 | 6 | 204 |
| 1h | Origin selectivity index of reach-related neurons | 5 | 7 | 229 |
| 2c | Spatial tuning of reach-related neurons | 7 | 8 | 252 |
| 2d | Target selectivity index of reach-related neurons | 7 | 8 | 252 |
| 2e | Targets eliciting response | 7 | 8 | 252 |
| 2f | K-means clustering | 7 | 8 | 252 |
| 2g | Seven-target reaching response profiles | 7 | 8 | 252 |
| 2h | Preferred target distribution | 7 | 8 | 252 |
| 3e | Neural state trajectories | 10 | 27 | N/A |
| 3f | Angle between neural state trajectories | 10 | 27 | N/A |
| 4c-e | Neural state trajectories during waterspout jump | 1 | 1 | N/A |
| 4g-h | Neural state trajectory distances | 3 | 4 | N/A |
| SF2g | Neuron lateralization of reach-related neurons | 14 | 44 | 1226 |
| SF3c | Hand tuning of reach-related neurons | 5 | 7 | 229 |
| SF3c | Hand eliciting response | 5 | 7 | 229 |
| SF5b | K-means clustering Silhouettes | 7 | 8 | 252 |
| SF5c | Spatial tuning of reach-related neurons in naïve mice | 4 | 4 | 189 |
| SF5d | Target selectivity index of reach-related neurons in naïve mice | 4 | 4 | 189 |
| SF5e | Three-target reaching response profiles in naïve mice | 4 | 4 | 189 |
| SF6a | Task-event triggered Calcium traces | 14 | 44 | 2863 |
| SF7b | Task-related neurons | 14 | 44 | 2247 |
| SF8c | Neural state trajectories – example session | 1 | 1 | 434 |
| SF8d | Explained variance by principal components | 10 | 27 | N/A |

**Extended Data Table 2: statistics**

| **Figure** | **Test** | **p-Value** | **Post-hoc** | **Comparisons** | **p-Value** |
| --- | --- | --- | --- | --- | --- |
| 1d bottom | Friedman | 7.93E-19 | Wilcoxon signed rank test | 1 vs 2  1 vs 3  2 vs 3 | 7.6048e-09  7.5871e-09  1.1453e-07 |
| 1f | Friedman | 3.6332e-04 | Wilcoxon signed rank test | Ori A vs Ori B (Left)  Ori A vs Ori B (Center)  Ori A vs Ori B (Right) | 0.0156  0.0156  0.0156 |
| 2e | Friedman | 1.71E-08 | Multiple comparisons rank-sums test with Bonferroni correction | 1 vs 6 1 vs 7 2 vs 5 2 vs 6 2 vs 7 3 vs 6 3 vs 7 | 0.0074 0.0074 0.0116 0.0002 0.0002 0.0023 0.0023 |
| 2h | Chi-Square | 0.0057 | N/A | N/A | N/A |
| SF2g | Friedman | 2.02E-07 | Wilcoxon signed rank test | Left vs Center  Left vs Right  Center vs Right | 7.6342e-04  3.2072e-06  0.4711 |
| SF3d | RM ANOVA | 0.0.0026 | Tukey-Kramer post-hoc | Left vs Both  Right vs Both | 0.011238  0.0066629 |
